# Time-course RNASeq of *Camponotus floridanus* forager and nurse ant brains indicate links between plasticity in the biological clock and behavioral division of labor

**DOI:** 10.1101/2021.03.27.433505

**Authors:** Biplabendu Das, Charissa de Bekker

**Affiliations:** Department of Biology, College of Sciences, University of Central Florida, Orlando, Florida, 32816; Genomics and Bioinformatics Cluster, University of Central Florida, Orlando, Florida, 32816

**Keywords:** carpenter ants, behavioral division of labor, circadian rhythms, ultradian rhythms, time-course RNASeq

## Abstract

**Background:** Circadian clocks allow organisms to anticipate daily fluctuations in their environment by driving rhythms in physiology and behavior. Inter-organismal differences in daily rhythms, called chronotypes, exist and can shift with age. In ants, age, caste-related behavior and chronotype appear to be linked. “Around-the-clock” active nurse ants are usually younger and, with age, transition into rhythmically active foragers. Moreover, ants can shift between these behavioral castes depending on social context. We investigated how changes in daily gene expression could be contributing to such behavioral plasticity in *Camponotus floridanus* carpenter ants by combining time-course behavioral assays and RNA-Sequencing of forager and nurse brains.

**Results:** We found that nurse brains have three times fewer 24h oscillating genes than foragers. However, several hundred genes that oscillated every 24h in forager brains showed robust 8h oscillations in nurses, including the core clock genes *Period* and *Shaggy*. These differentially rhythmic genes consisted of several components of the circadian entrainment pathway, and showed enrichments for functions related to metabolism, cellular communication and protein modification. We additionally found that *Vitellogenin*, known to regulate division of labor in social insects, showed robust 24h oscillations in nurse brains but not in foragers. Furthermore, the protein products of several genes that were differentially expressed between the two ant castes were previously found in the trophallactic fluid of *C. floridanus*. This suggests a putative role for trophallaxis in regulating behavioral division of labor through caste-specific gene expression.

**Conclusion:** We provide a first look at the chronobiological differences in gene expression between forager and nurse ant brains. This endeavor allowed us to identify putative molecular mechanisms underlying plastic timekeeping. Several components of the ant circadian clock and its output can seemingly oscillate at different harmonics of the circadian rhythm. We propose that such chronobiological plasticity has evolved to allow for distinct regulatory networks that underlie behavioral castes, while supporting swift caste transitions in response to colony demands. Behavioral division of labor is common among social insects. The links between chronobiological and behavioral plasticity that we found in *C. floridanus*, thus, likely represent a more general phenomenon that warrants further investigation.

## Background

Living organisms exhibit adaptive rhythms in physiology and behavior as a way to anticipate predictable daily fluctuations in their environment [1–3]. Such daily rhythms are ubiquitous and have been discovered in both unicellular and multicellular organisms [4–9], including eusocial Hymenopterans such as ants and bees [10–16]. These rhythms are driven by endogenous molecular feedback loops that are capable of entraining to external time cues, known as Zeitgebers, which can be both abiotic (e.g., light and temperature cycles) and biotic (e.g., presence of food and predators) [17–20]. In the majority of model organisms studied thus far, light appears to be the strongest Zeitgeber [19, 21]. However, it has been suggested that in Hymenopterans with complex social behaviors, temperature cues and social environment could be more potent Zeitgebers than light [22–26]. Though, a more thorough molecular understanding of the Hymenopteran clock and its role in the social organization of insect colonies is needed to confirm this.

The limited knowledge that we currently have of the Hymenopteran clock stems from a handful of studies done with the honeybee *Apis mellifera* and a few ant species including the carpenter ant *Camponotus floridanus* [14–16, 27–29]. This is in stark contrast with our vast molecular understanding of the circadian clock of *Drosophila melanogaster*, which has been extensively studied and is often used as a reference model for insect circadian clocks in general (reviewed in [30–32]). At the cellular level, the circadian clock consists of an autoregulatory transcription-translation feedback loop (TTFL) that requires around (circa) 24 hours (dia) to complete one cycle. The circadian TTFL is considered to be an ancient timekeeping mechanism conserved in plants, fungi and animals [2, 30, 33]. In the insect model organism *Drosophila*, the TTFL consists of the activator complex CLOCK-CYCLE (BMAL1-CLOCK in mammals) that binds to and activates transcription of the repressor gene *Period* (*Per*). Upon translation in the cytoplasm, PER heterodimerizes with TIMELESS (CRYPTOCHROME in mammals), translocates into the nucleus and inhibits the CLK-CYC activator complex, thus closing the feedback loop [34, 35]. This loop is further coupled with multiple auxiliary phosphorylation-dephosphorylation cycles, that are necessary for a functional 24-hour clock [34, 35]. Several kinases (e.g., Shaggy, Double-time, Nemo, Casein Kinase-2 and Protein Kinase A) and phosphatases (e.g., Protein phosphatase 1 and Protein phosphatase 2A) involved in such auxiliary cycles have been discovered in *Drosophila* (reviewed in [31]). Once entrained, the circadian clock drives daily oscillations in gene expression and protein production that in turn bring about rhythms in physiology (e.g., metabolism and immune function) and behavior (e.g., locomotion and feeding) [36].

In addition to being endogenous and entrainable, circadian clocks are also inherently plastic; the phase, amplitude and period length with which circadian processes oscillate can change with an organism’s age or social environment [37–42]. Such changes give rise to phenotypes that differ in their exact timing of activity onset relative to sunset or sunrise, known as “chronotypes” [43–46]. Social insects, which exhibit complex social organization and a decentralized division of colony labor, provide a striking example of plastic chronotypes which appear to be tightly associated with an individual’s behavioral role or caste identity within the colony [47, 48]. In ants and bees, broadly two distinct behavioral castes emerge from division of colony labor among non-reproductive “workers”: 1) extranidal foragers that primarily gather food in an environment with daily cycling abiotic conditions and 2) intranidal nurses that perform brood care within a nest with little to no abiotic fluctuations [49]. In most species studied so far, forager ants and bees show robust daily rhythms in locomotion and extranidal visits whereas nurses display “around-the-clock” activity patterns within the dark nest chambers [26, 47, 48, 50]. The presence or absence of circadian locomotory rhythms, thus, appear to be caste-associated. Seemingly, these rhythms are also plastic since foragers coerced into tending brood will begin to show “around-the-clock” activity whereas brood-tending nurses develop robust locomotory rhythms upon removal from the colony [15, 25, 27, 51]. For example, in the carpenter ant *Camponotus rufipes*, nurses showed a rapid development of rhythmic activity patterns when isolated from the colony and placed under cycling light-dark conditions [48]. This rhythmic activity persisted under constant darkness conditions in the absence of brood [48]. Similarly, isolated individuals of the ant species *Diacamma indicum*, showed rhythmic activity under LD cycles in the absence of eggs and larvae, but transitioned to nurse-like “around-the-clock” activity in their presence [25]. As such, circadian rhythms in locomotory behavior appear to be regulated by an individual’s social context and behavioral role in the colony [25, 26, 48, 52]. This is in line with the finding that social cues, such as colony odor or substrate-borne vibrations, can be potent Zeitgebers in social insects and can even override photic entrainment [23, 24].

The molecular aspects of plastic timekeeping and its role in driving behavioral plasticity that gives rise to colony-wide division of labor in ants, and other social insects, are largely unexplored. Exposing the mechanisms of plastic timekeeping in ants, and how they connect to behavioral phenotypes, could be essential in our understanding of eusocial behavior and regulation of colony functioning. A first step in this direction has been made by Rodrigues-Zas and colleagues, who investigated circadian gene expression in honeybee forager and nurse brains through a time-course microarray study [16]. However, this study identified only 4% of all protein coding genes as rhythmic, which seems almost certainly a vast under-representation considering the abundance of clock-controlled genes that have been found in other organisms [53–59]. No other genome-wide reports that assess daily rhythms in gene expression seem to exist for Hymenoptera despite the availability of newer high-throughput sequencing techniques and improved rhythm detection software [60, 61]. As such, a major knowledge gap regarding the inner workings of social insect clocks, and especially those of ants, remain. This greatly limits our ability to investigate how biological clocks could be interacting with social cues to produce functionally distinctive behavioral castes with their own characteristic chronotypes.

Our current study aims to address this knowledge gap by investigating rhythmic gene expression, throughout a 24h-day, in brains of *Camponotus floridanus* nurse and forager ants. The Florida carpenter ant, *C. floridanus*, produces large colonies with several thousand workers, organized in both behavioral and morphological castes. This species is considered an urban pest [62] and is frequently used in a wide variety of social insect studies (e.g., [63–72]). To collect forager and nurse ants of *C. floridanus*, we conducted a time-course experiment in a complex, large colony setup that allowed us to quantify circadian foraging behavior of the colony and identify individuals based on their behavioral caste. We subsequently used the brains of collected foragers and nurses for RNASeq to fulfill three primary objectives: (1) to investigate the extent of rhythmic gene expression for both castes, (2) to characterize the similarities and differences in their daily transcriptomes, and (3) to identify putative mechanisms that could allow nurse ants to possess a functional timekeeping machinery despite no apparent circadian rhythms in daily activity. We found that nurse brains harbored a reduced number of circadian genes as compared to foragers. Yet, we discovered that several genes with robust circadian expression in forager brains were not entirely arrhythmic in nurses. Rather, these genes oscillated with 12-hour and 8-hour periodicities (the core clock gene *Period* being one of them). We discuss the possibility that such plasticity in clock and clock-controlled gene expression could facilitate swift nurse to forager transitions and vice-versa. Furthermore, we used functional enrichments of gene ontology annotations to identify biological processes that are seemingly under clock-control in *C. floridanus* brains, and highlight the ones enriched for genes that cycled at different periodicities in the two ant castes. Additionally, we report on genes that were expressed at vastly different levels in the brains of the two ant castes, throughout the day. The protein products of several of these differentially expressed genes have been discovered in the trophallactic fluid of *C. floridanus* [69, 73]. As such, we discuss the possibility that division of labor and the regulation of behavioral chronotypes in ant societies is trophallaxis-mediated.

## Results and Discussion

### Daily rhythms in colony behavior of *Camponotus floridanus*

*Camponotus floridanus* is known to be largely nocturnal both in nature (personal field observations, [62]) and in the lab [67, 68]. Despite this knowledge, we first had to entrain and quantify the colony-level behavioral rhythms of *C. floridanus* to be able to reliably investigate the daily gene expression underlying their seemingly clock-regulated behavioral activity. Therefore, we recorded extranidal visits of a large *C. floridanus* colony, housed in a darkened nest, that we attached to a foraging arena subjected to a 12h:12h LD cycle (see Methods section for more details). Subsequently, we counted the number of foraging ants throughout the day that were actively feeding or present on the feeding stage (Fig. 1, “FS” or feeding activity) as well as in the remainder of the foraging arena (Fig. 1, “FA” or general foraging activity). We defined the colony’s total foraging activity (Fig. 1, “Total activity”) as the sum of FS and FA at any given time. The first signs of initial colony entrainment were visible through the early establishment of a day-night rhythm in FA (Fig. 1, Day 1-5). In the following 3 days, we performed mark-and-recapture to identify ants of the foraging caste. During this time the FA rhythm was somewhat less pronounced but managed to stay intact (Fig. 1, Day 6-8). From Day 9 onwards, both FS and FA showed pronounced day-night rhythms that persisted during and beyond the sampling day (Fig. 1, Day 9-15). These day-night rhythms followed a consistent pattern with increased foraging activity during the night-time as compared to the daytime, similar to previously reported locomotory rhythms of isolated *C. floridanus* ants [68]. Thus, based on extranidal activity of the foraging caste, the colony established robust nocturnal activity rhythms as it would in nature by entraining to the light Zeitgeber we provided.

**Figure 1.**
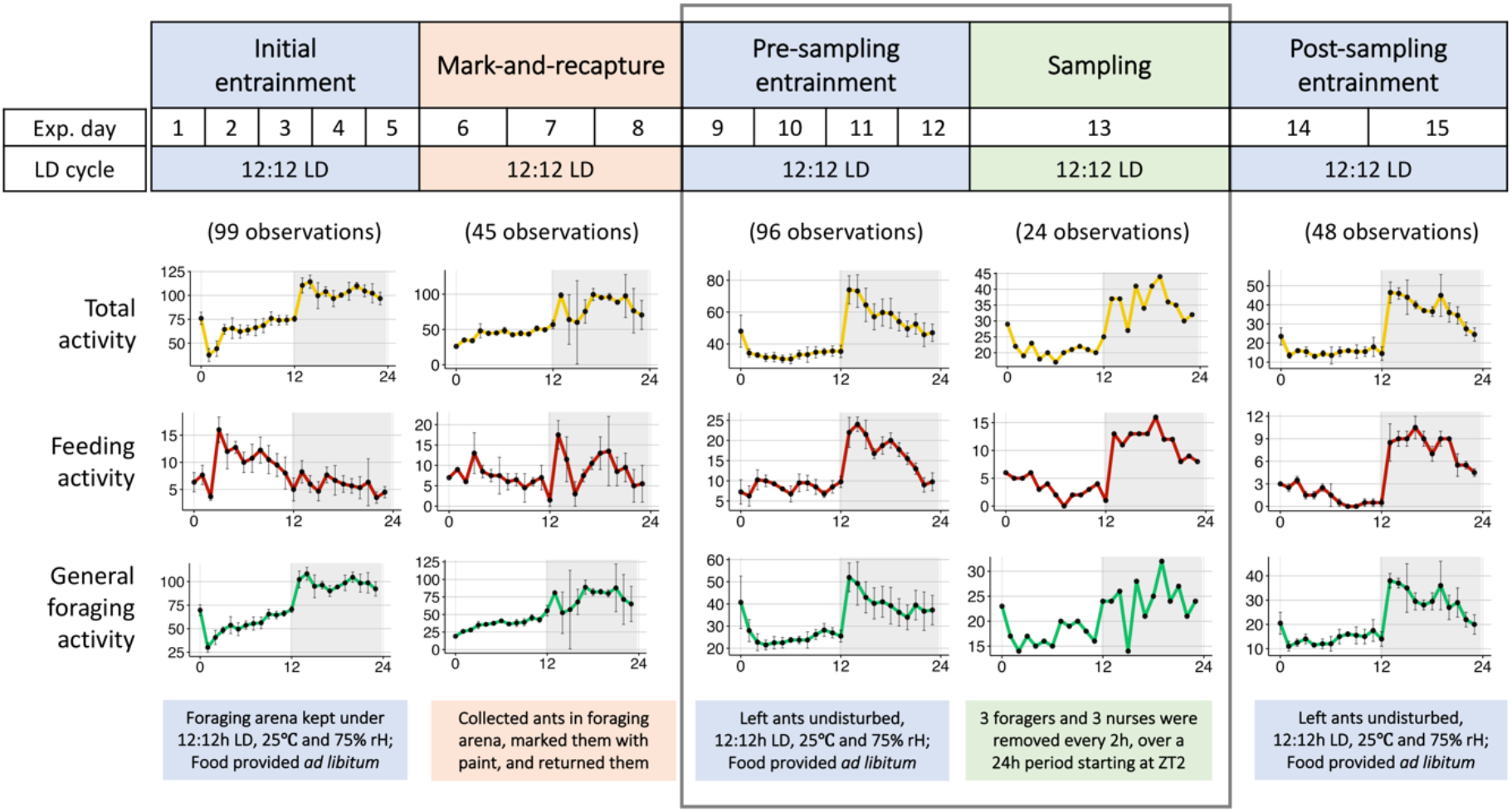
Daily rhythms in colony activity. The top panel shows the experimental timeline and the bottom graphs show the mean (±SE) daily extranidal activity of the ant colony during each phase of the experiment. During the entire experiment, the foraging arena was kept at 25°C, 70% rH and under oscillating 12h:12h light-dark (LD) cycles. Undisturbed phases under light-dark cycles are shown in blue, while experimental phases of disturbance are shown in orange (mark-and-recapture of foragers) and green (sampling of ants for RNASeq). For each plot, colored lines connecting the dots represent average activity while black bars represent one standard error around the mean. The y-axis represents number of ants and the x-axis represents Zeitgeber Time (ZT) during the 12h:12h LD cycle. The shaded part of the plots represents the dark phase (ZT12-24). The number of ants actively feeding or present on the feeding stage is plotted as the feeding activity (FS). The general foraging activity (FA) is the number of ants present in the foraging arena but not on the feeding stage. The total activity is the sum of FA and FS, representing the total extranidal activity of the colony at a given time. The number of observations used to calculate the mean (± SE) activity for each phase are shown in parenthesis at the top of the plots. Missing data points during ‘Initial entrainment’ and ‘Mark-and-recapture’ were due to inability to get accurate count of ants from video frames and a recording failure, respectively.

To further characterize the behavioral rhythms in the entrained *C. floridanus* colony and to investigate the potential behavioral effects of the disturbance introduced by the mark-recapture, we performed wavelet analyses [74] on the foraging data collected during the four-day period just after mark-recapture and prior to sampling (Fig. 1, Day 9-12). *Camponotus floridanus* ants of the foraging caste showed significant circadian rhythms in FS and FA (Fig. 2A). Average wavelet powers indicated that both FS and FA activity profiles comprised of significant waveforms with a period length close to 24 hours (Fig 2A). Neither FS nor FA activity peaked exactly at lights-off (ZT12). Rather, we noticed a sharp increase in both activities about an hour later (∼ZT13) (Fig 1, Day 9-12). After peaking around ZT13-15, both FS and FA activity continued to decrease throughout the night and reached their daily minimum shortly after lights were turned on (ZT2-4) (Fig 1, Day 9-12). Central Florida (the location of colony collection) receives an average of 12 ± 1 (mean ± SE) hours of sunlight per day, and dusk lasts for 84 (± 5) minutes after sunset (Additional File 1A, data retrieved from www.timeanddate.com). In our experimental setup, we chose an abrupt light-dark transition, and hence, did not provide twilight cues. Therefore, the stark increase in extranidal activity within an hour post lights-off, could be indicating an endogenous dusk-entrainment in colony foraging activity. Taken together, the colony activity rhythms that we observed for *C. floridanus* – primarily circadian, and predominantly nocturnal, with a dusk-phase – largely resembled previously reported activity patterns [68]. This indicates that the experimental setup that we designed allowed us to collect daily gene expression data related to expected ant daily activity patterns.

**Figure 2.**
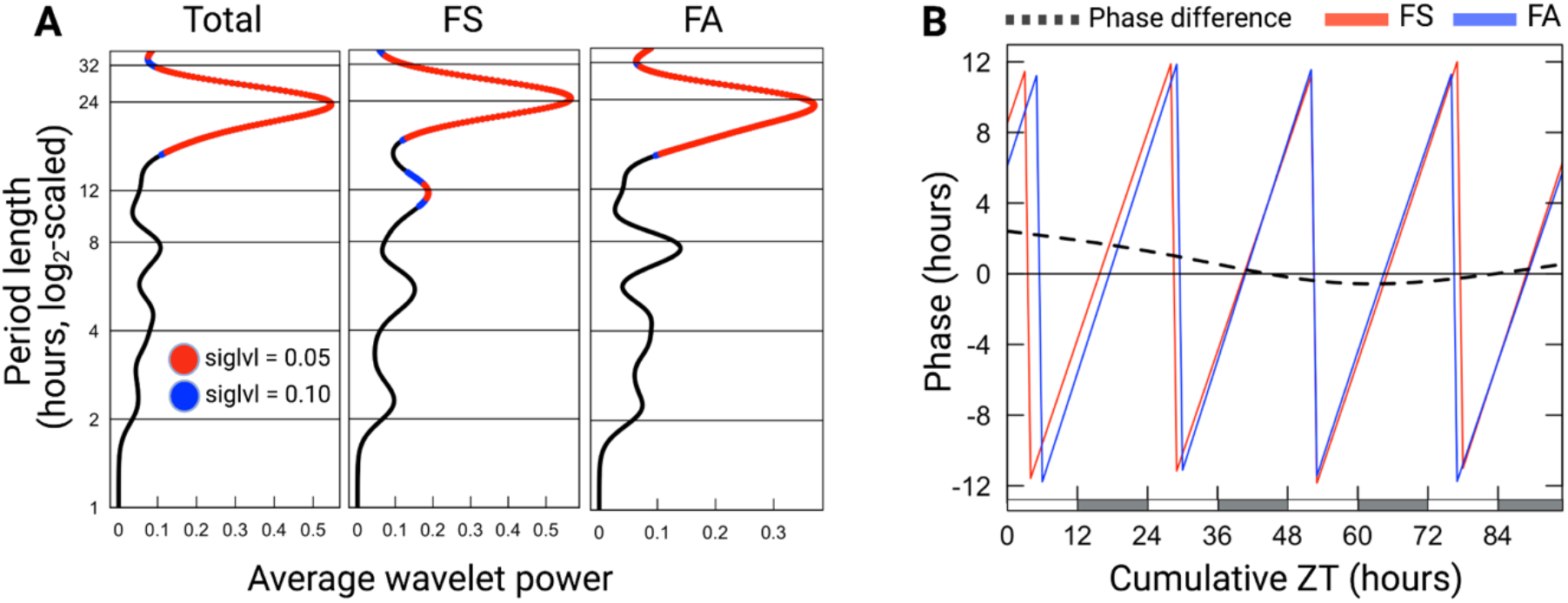
Wavelet analysis of feeding and general-foraging activity rhythms. **(A)** Dominant periods identified using wavelet decomposition of each activity profile during the continued entrainment phase (Day 9-12). The x-axis shows the average wavelet power for different period lengths. The y-axis shows the period length (log2-scaled) in hours. Significant period lengths (siglvl < 0.05) are shown in red, and the peak indicates the dominant period having the most power (around 24h for all three activity profiles, and an additional 12h peak observed in feeding bouts); **(B)** The plot shows the phase (on the y-axis) of feeding bouts (FS) and general foraging activity (FA) during continued entrainment. The dotted line indicates the phase difference of FS over FA during the same time period. Positive phase difference indicates that FS leads FA. The x-axis shows the cumulative hours passed since disturbance due to mark-and-recapture (Cumulative ZT).

In addition to the dominant circadian rhythm, we detected a significant circa-12h rhythm in FS (Fig. 2A). Inspection of FS power-spectra over the four days of continued-entrainment revealed that, while the circadian rhythm was sustained throughout, the 12h rhythm was only significantly present during the first 36 hours post disturbance. Within this 36h time-period, integration of the 12h and 24h FS waveform improved fit (Additional File 1B). A possible explanation for the presence of this short-lived 12h activity rhythm could be that it played a role in catching up with feeding needs of the colony in the initial hours after disturbance. The removal of foragers during mark-recapture most likely desynchronized the colony’s daily feeding pattern and might explain the lack of a clear circadian activity in FS and a diminished overall 24h foraging pattern during the mark-recapture period (Fig 1; Day 6-8). As such, we enquired if the circa-12h rhythm in FS could be important to re-establish a rhythmic colony feeding behavior that is synchronous to the colony’s general foraging activity. To this end, we calculated the phase difference of the 24h-wavelets for FS-over-FA throughout the four days post mark-recapture (Fig. 2B). At the start of pre-sampling entrainment (i.e., right after disturbance by mark-recapture), FS was found to lead FA by more than two hours. Approximately 36 hours into the pre-sampling entrainment period, the phase difference reduced to zero; 24h-rhythms in FS and FA aligned. Subsequently, the phase difference between FS and FA remained close to zero (Fig. 2B). This data suggests that, indeed, after three consecutive nights of disrupted feeding, the colony attempted to get back on track through a short initial phase shift between FS and FA. Once synchrony between the phases of the two activities was restored, it was maintained. The intermittent 12h feeding peaks observed during the first 36h after mark-recapture (Additional File 1B) likely contributed to restoring this synchrony.

### General patterns of gene expression in *C. floridanus* brain tissue

After twelve days of LD entrainment, we collected three *C. floridanus* foragers and nurses from the colony every 2 hours, over a 24-hour period (Fig. 1, Day 13). Individuals that were collected in the foraging arena and paint-marked as part of our mark-recapture efforts were collected as foragers. Unmarked individuals that interacted with the brood inside the dark nest chambers were collected as nurses. We subsequently used RNA-Seq to obtain the transcriptome profiles of forager and nurse brain tissue. Of the 13,808 protein coding genes annotated in the *C. floridanus* genome [70], 8% were not expressed (i.e., FPKM = 0) in any of the samples collected (Additional File 2, sheet 1). These 1130 non-expressed genes were enriched in multiple biological processes: DNA integration, DNA replication, telomere maintenance, proteolysis and apoptotic process, and several molecular functions including hormone activity (Additional File 2, sheet 2). More than half of all genes annotated to be involved in nucleotide binding (56% of 27 genes), DNA integration (55% of 86), and those located in the extracellular matrix (53% of 38) did not exhibit any expression in *C. floridanus* brains.

Furthermore, 19% of the *C. floridanus* genes (2640 genes) were only lowly expressed in forager and nurse brains (i.e., 0 < FPKM ≤ 1) throughout the day (Additional File 2, sheet 1). The majority of genes involved in olfactory and gustatory functions in *C. floridanus* were among these lowly expressed genes (93% of 363 genes involved in sensory perception of smell and 73% of 26 genes involved in sensory perception of taste) (Additional File 2, sheet 2). Notably, majority of the genes involved in hormone activity (69% of 16), metallopeptidase activity (86% of 110), and nucleotide binding (85% of 27) were found to be enriched among the genes that showed either no or low expression (Additional File 2, sheet 2). The clear overrepresentation of certain gene functions among genes that were either lowly or not expressed necessitated the use of a reduced background gene set for subsequent enrichment analyses that consists of only those genes that were actually expressed. This, to avoid obtaining gene function enrichments that merely reflect brain tissue specific gene expression. We classified genes to be expressed in *C. floridanus* brains if mRNA levels were greater than 1 FPKM for at least one time point, for either behavioral caste, during the 24h sampling period.

We found 71% (i.e., 9843 genes in foragers and 9872 genes in nurses, Additional File 2, sheet 3) of all protein coding genes to be expressed in ant brains. Of these genes, 166 were uniquely expressed in the forager brains and 195 in nurses. One *odorant receptor 4-like*, two *odorant receptor 13a-like*, and two other uncharacterized odorant receptor genes were among those uniquely expressed in forager brains, along with several proteases. In addition to significant enrichments in olfaction and proteolysis-related biological processes, uniquely expressed genes in foragers were also enriched in the cellular component nucleosome and included several histone-related genes (Additional File 2, sheet 4). In comparison, genes uniquely expressed in nurses were enriched in redox and lipid metabolic processes and included several putative *cytochrome P450* and *lipase 3-like* genes (Additional File 2, sheet 4). This is in line with the canonical behavioral and physiological differences that characterize foragers and nurses in a social insect colony. A fine-tuned olfactory and gustatory repertoire in foragers is essential for trail-following and other general foraging tasks. In contrast, metabolic processes have been previously found to be upregulated in intranidal nurse workers that are usually tasked with larval feeding and brood care [75]. This indicates that the expression data that we obtained is likely a good representation of the gene expression profiles that are characteristic for both castes.

### Circadian rhythms in gene expression

We used the non-parametric algorithm empirical JTK Cycle (eJTK) [76, 77] to detect circadian gene expression patterns in forager and nurse ant brains. Of the 10,038 genes expressed in *C. floridanus* brains, 42% (i.e., 4242 genes) had significant circadian expression patterns in either foragers or nurses (Additional File 3, sheet 1 and 2). The number of putative circadian genes in foragers was almost three times higher (i.e., 3569 genes; Fig 3A and B, indicated with “for-24h”) as compared to nurses (i.e., 1367 genes; Fig 3A and C, indicated with “nur-24h”). Only 16% of all identified circadian genes cycled in both behavioral castes with a 24h rhythm (i.e., 694 genes; Fig 3A and D, indicated with “for-24h-nur-24h”), which represents half of all the circadian genes that we identified in nurses. The reduced number of circadian genes in nurses is consistent with the previous time-course microarray study done in honeybees (541 probes in forager bees and 160 probes in nurse bees were found to be circadian) [16]. This suggests that a reduced circadian control at the level of gene expression in “around-the-clock” active nurses as compared to rhythmically active foragers likely persists across social Hymenoptera that display division of labor.

**Figure 3.**
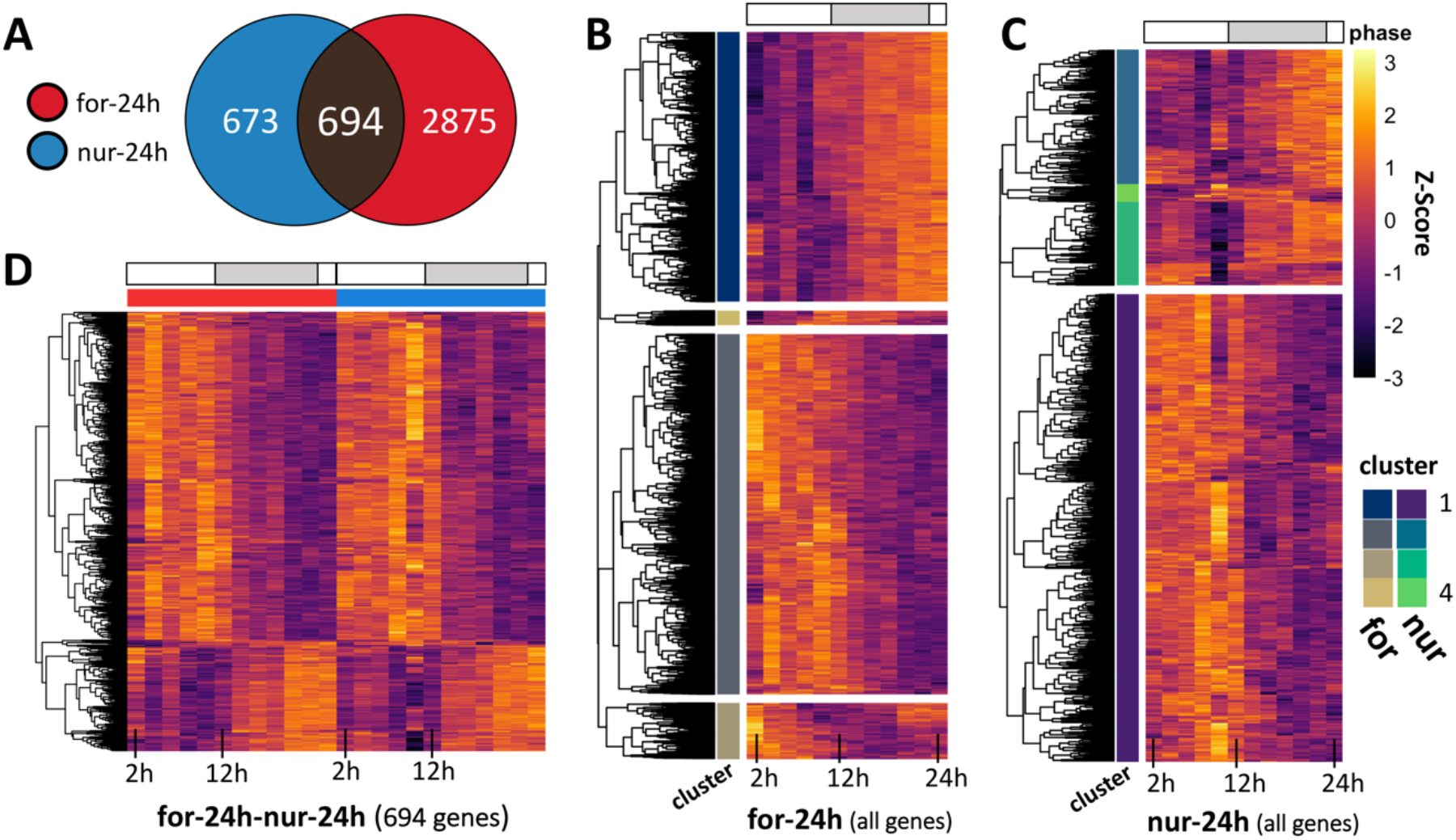
Circadian rhythms of gene expression in the ant brain. (**A**) Venn-diagram showing the number of genes significantly oscillating every 24h in forager (for-24h) and nurse (nur-24h) brains. The heatmaps show the daily expression (z-score) patterns of all identified 24h-oscillating genes in **(B)** foragers (for-24h), **(C)** nurses (nur-24h), and **(D)** both foragers and nurses (for-24h-nur-24h). Each row represents a single gene and each column represents the Zeitgeber Time (ZT) at which the sample was collected, shown in chronological order from left to right (from ZT2 to ZT24, every 2h). The grey bar above the heatmaps runs from ZT12 to ZT24 and indicates the time during the light-dark cycle in which lights were off. Both for-24h and nur-24h genes were hierarchically clustered into four clusters. The cluster identity of each gene is indicated in the cluster annotation column.

After identifying putative 24h cycling genes in the two behavioral groups, we asked if they contained functional annotations with coordinated temporal peak activity (i.e., are certain biological functions “day-peaking” or “night-peaking”) and if such a temporal division of clock-controlled processes can be found in both foragers and nurses. To answer these questions, we used an agglomerative hierarchical clustering framework to group the circadian genes in foragers and nurses into four gene clusters (Additional File 3, sheet 3 and 4). We followed this analysis by identifying significantly enriched gene ontology (GO) terms for each identified gene cluster.

The choice of four clusters was aimed to demarcate, if possible, potential day-, night-, dawn-, and dusk-peaking genes. Using this method, we identified that more than half of all circadian genes in foragers showed a peak activity during early-to-mid daytime (1916 genes, Fig. 3B, for-24h_Cluster2). The majority of the remaining genes showed peak expression activity around late night-time (1417 genes, Fig. 3B, for-24h_Cluster1). Additionally, one of the two smaller clusters of genes that cycled with a 24h rhythm in foragers (74 genes, Fig. 3B, for-24h_Cluster4) appeared to peak at dusk with an acrophase around ZT12-14. Among these dusk-peaking genes we identified the putative insect melatonin receptor *trapped in endoderm* (*tre1; MTNR1a* in mammals), which has been reported to be central to the dusk/dawn entrainment pathway in humans (Table 1) [78–80]. The genes in nurse brains that showed 24h rhythms also primarily clustered into two groups – day-peaking (909 genes, Fig 3C, nur-24h_Cluster1) and night-peaking genes (261 genes, Fig 3C, nur-24h_Cluster2) – with only a few genes in the remaining two clusters (Cluster 3, 162 genes; Cluster 4, 35 genes).

**Table 1.**
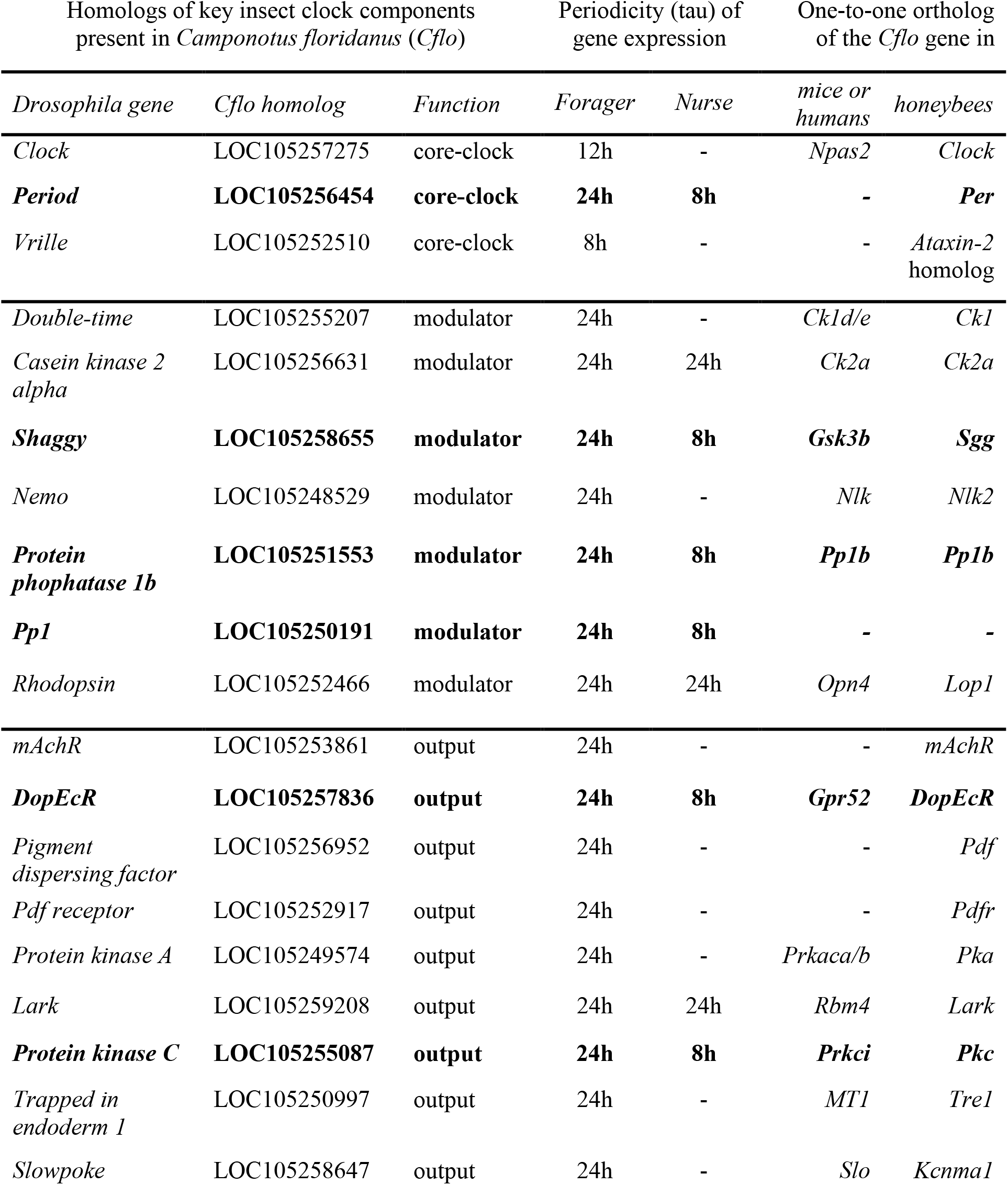
Clock components of Camponotus floridanus and their gene expression patterns in forager and nurse brains. . The table below lists the *C. floridanus* homologs of several *Drosophila* core-clock, clock-modulator and clock-output genes. The periodicity (tau) of rhythmic gene expression in the brain, if any, is indicated for both foragers and nurses. The one-to-one ortholog of the identified *C. floridanus* gene in mammals and honeybees is also provided. A dash in the periodicity column indicates that no significant daily rhythms were detected for the *C. floridanus* gene, whereas a dash in the ortholog columns indicates that no one-to-one orthologs of the *C. floridanus* gene was detected. The genes that show differential rhythmicity, oscillating at two distinct periodicities, in the two ant castes are shown in bold.

Despite the relatively smaller number of day-peaking and night-peaking circadian genes in nurses, we found functional enrichments comparable to those found in foragers. The night-peaking gene clusters in foragers and nurses were both enriched in genes with the annotated GO terms: regulation of transcription (DNA-templated), signal transduction and protein phosphorylation (Additional File 3, sheet 5). This indicates that a significant number of night-peaking circadian genes in nurse and forager brains seem to be involved in cell-cell communication, gene expression, and protein modification. The day-peaking circadian gene clusters in both behavioral groups were enriched for genes involved in metabolism (glycosylphosphatidylinositol (GPI) anchor biosynthesis) (Additional File 3, sheet 5). In addition, the circadian gene clusters in foragers were enriched for multiple other biological processes that were not found to be enriched in nurses. The day-peaking genes in foragers were enriched for GO terms that concerned response to stress, as well as tRNA, mRNA and translational processes, and terms involved in post protein processing such as folding and transport (Additional File 3, sheet 5). Night-peaking genes in foragers were additionally enriched in terms such as regulation of transcription by RNA polymerase II, multicellular organism development, protein homooligomerization, microtubule-based movement, G protein-coupled receptor signaling pathway, and ion transmembrane transport (Additional File 3, sheet 5). This temporal segregation of clock-controlled processes in foragers appears to be in line with findings from previous studies done on the fungus *Neurospora crassa*, mammals and flies [55, 57, 81]. However, while the daily transcriptome of rhythmic foragers revealed the expected temporal separation, nurse gene expression showed a much more limited temporal organization. This provides further evidence for a reduced circadian control in “around-the-clock” active nurses as compared to rhythmically active foragers.

The question that remains is if the shared functional enrichments among the 24h rhythmic genes in both ant castes encompass the same exact genes or if they are different but with similar functions. To answer this question, we analyzed the functional annotations of the 694 circadian genes that were shared between foragers and nurses. Hierarchical clustering revealed that these genes predominantly peaked during the daytime (Fig. 3D) and that the shared day-peaking genes were significantly enriched in the functional annotation GPI anchor biosynthesis (genes *Pig-b, Pig-c, Pig-g, Pig-m,* and *Mppe*) (Additional File 3, sheet 5). However, the relatively smaller set of shared night-peaking circadian genes was not enriched in any functional annotations. This suggests that the night-peaking activity of regulation of transcription (DNA-templated), signal transduction and protein phosphorylation are mostly due to different sets of circadian genes in foragers and nurses, but with similar functions. In contrast, GPI anchor biosynthesis activity appears to be driven by the same day-peaking circadian genes in both ant castes.

The molecular underpinnings of timekeeping in nurse ants, and other animals with “around-the-clock” activity, is still elusive [14, 16, 82]. To find candidate genes presumably involved in daily timekeeping in *C. floridanus* nurses, we queried the circadian genes that they shared with foragers for known components of the insect clock (Additional File 4). The shared day-peaking gene cluster contained one known clock output gene (i.e., *Lark*) and two genes known to modulate the circadian clock – *Casein kinase 2 alpha* (*Ck2a*) and the light-dependent *Rhodopsin* (*Rh6*; orthologous to mammalian *Opn4*) (Table 1, Fig. 4). Along with other members of the opsin gene family, the *Rh6* gene in *Drosophila* has been shown to also have light-independent functions in thermosensation (in larvae) and hearing (in adults) [83, 84]. The auditory role of opsins, likely mediated by mechanotransduction [85], could be especially relevant for circadian entrainment in social insects. Ants and bees are known to use vibroacoustic means such as “drumming” behavior (i.e., vibrations produced by tapping the nest substrate with their head and gaster) to communicate within dark nest chambers [86–89]. Moreover, there is recent evidence that substrate-borne vibrations are potent social Zeitgebers capable of entraining the circadian clock of newly emerged honey bees housed in the dark [24]. These substrate-borne vibrations could potentially play a similar role in the social entrainment of nurse ants through the light-independent involvement of a rhodopsin-mediated mechanosensory pathway [85], while extranidal foragers might also make use of its light-dependent functions.

**Figure 4.**
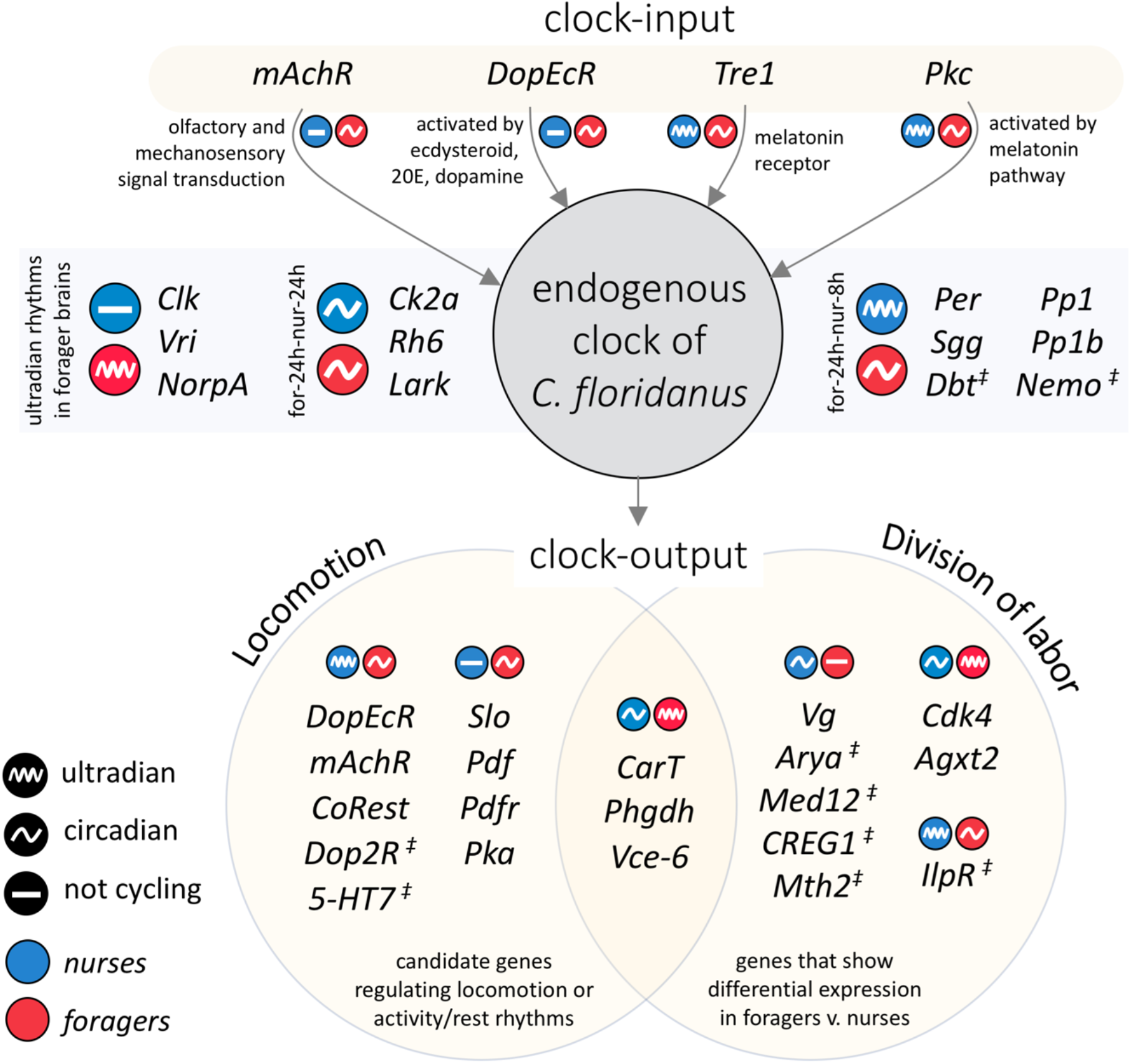
Potential links between chronobiological plasticity and behavioral division of labor in *C. floridanus*. The infographic shows differences in rhythmic expression in forager and nurse brains for several genes involved in entrainment of the endogenous clock (clock-input), proper functioning of the endogenous clock, and the clock-controlled pathways (clock-output) that likely regulate locomotion and division of labor in ants. The symbol “‡” indicates that gene expression for that gene shows a trend of rhythmic expression in one of the ant castes (Additional File 5) but was not significant (p ≥ 0.05). Ultradian rhythms include both 8h and 12h oscillations. The following genes has been abbreviated in the figure but not in the text: *Venom-carboxylesterase-6* (*Vce-6*), *Arylphorin-subunit-alpha* (*Arya*).

In addition to *Rh6,* the kinase *Ck2a* showed robust 24h rhythms and a near-perfect alignment in gene expression between the behavioral groups (Additional File 3, Fig. 4). *Ck2a* encodes the catalytic subunit of the circadian protein, Casein Kinase 2 (CK2). In *Drosophila*, CK2 appears to regulate rhythmic behavior by phosphorylating the core clock proteins PERIOD (PER) and TIMELESS (TIM) [90–93]. This CK2-mediated phosphorylation is perceived as a rate-limiting step in the circadian clock, important for a functional 24h transcription-translation feedback loop [93]. The central role of CK2 in regulating the endogenous clock in other organisms suggests a potential role of *Ck2a* in maintaining a functional 24h oscillator in both, “around-the-clock” active nurses and rhythmically active foragers. However, other homologs of genes encoding core clock proteins, such as PER, were not present among the circadian genes that were shared between foragers and nurses (Table 1, Additional File 3).

### Ultradian rhythms in gene expression

“Ultradian rhythms” in gene expression refer to significantly oscillating expression patterns around the second and third harmonic of circadian rhythms (i.e., genes cycling with periodicities of 12 hours and 8 hours, respectively). Such rhythms can be found in a wide range of species [94–100], and examples in which organisms switch from circadian to ultradian gene expression due to changes in environmental circumstances have been reported [101]. When we visually inspected the expression of several genes that exhibited circadian rhythmicity in foragers but not in nurses, we noticed that the expression of multiple such genes in nurses was relatively dampened but seemed to oscillate at a frequency higher than 24 hours. As such, we used eJTK to detect if any genes were expressed with significant ultradian rhythms (Additional File 6). We identified a comparable number of genes that cycled with a 12h period in forager and nurse brains (i.e., 148 and 193, respectively), and 2 genes that showed 12h period in both castes (Fig. 5A). In foragers, the core-clock gene *Clock* (*Clk*) was present among the 12h oscillating genes (Table 1, Fig. 4). However, we did not detect circadian or ultradian rhythmicity in *Clk* expression in nurses (Table 1). As for genes that oscillated with a robust 8h rhythm, we discovered 229 such genes in forager brains and about twice as many (550 genes) in nurses. Only three genes showed an 8h cycling pattern in both behavioral castes (Fig 5A).

**Figure 5.**
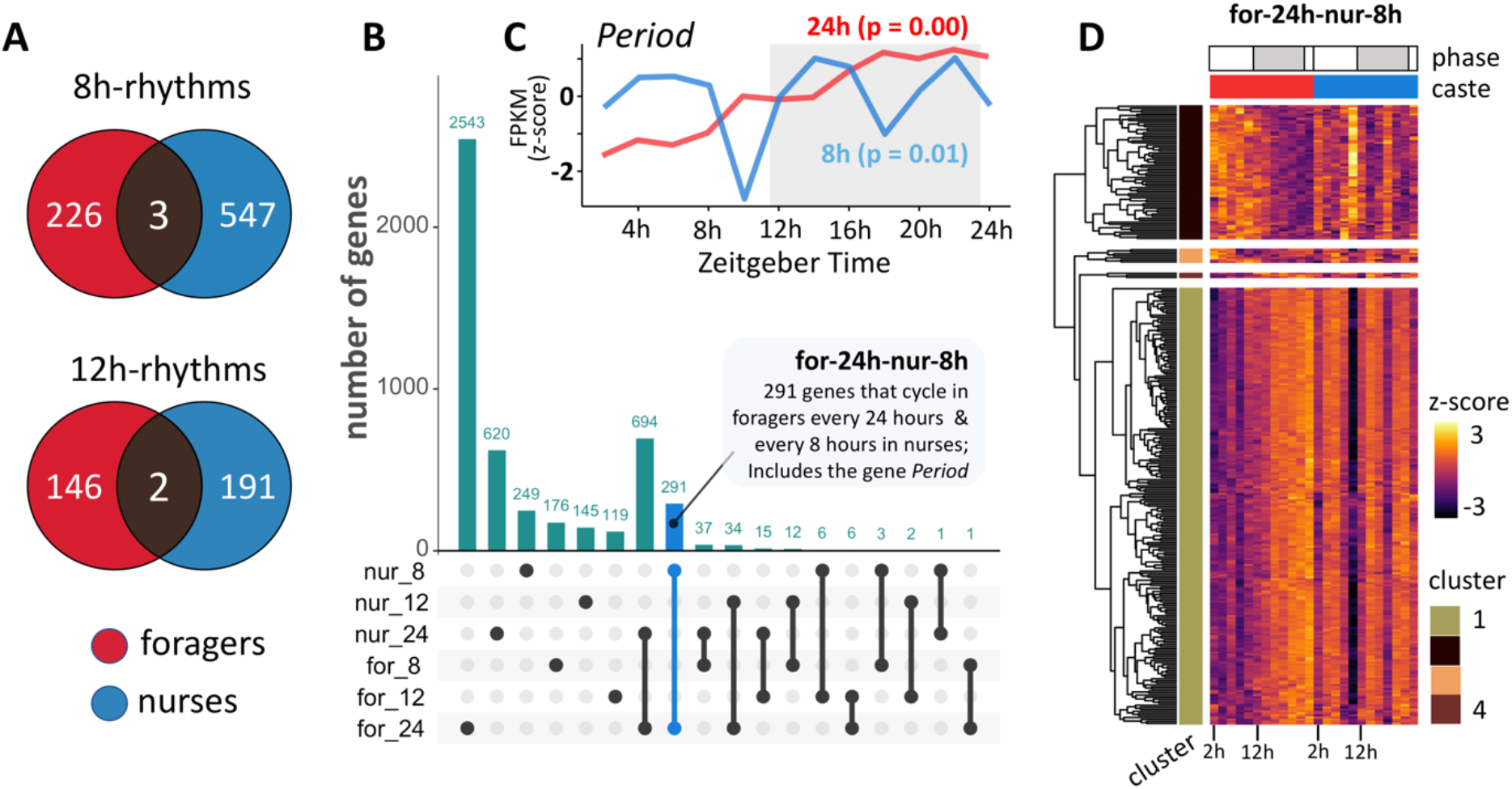
Ultradian rhythms and caste-associated differential rhythmicity in gene expression. (**A**) Venn-diagrams showing the number of genes with significant ultradian expression in the ant brain, oscillating every 8-hour (8h-rhythms) and 12-hour (12h-rhythms); (**B**) Upset plot showing the number of genes uniquely expressed in, and shared between, circadian (24h) and ultradian (8h and 12h) gene sets. Each bar represents a unique intersection between the six circadian and ultradian genesets (e.g., for_24: 24h-oscillating genes in foragers, nur-12: 12h-oscillating genes in nurses, etc.). A gene is binned only once, and as such, belongs to only one intersection. Dark circles indicate the gene sets that are part of a particular intersection. For example, the first circle indicates that there are 2543 genes that are uniquely cycling in foragers with a 24h period (for_24). Similarly, the blue bar indicates that there are 291 genes that have a significantly circadian expression in foragers but cycle every 8-hours in nurses (for-24h-nur-8h); **(C)** Caste-associated differential rhythmicity in the expression of the core clock gene *Period* is shown. The expression of *Per* cycles every 24-hours in forager brains (red) and every 8-hours in nurses (blue); p-values obtained from eJTK are provided in parenthesis. The Zeitgeber Time is indicated on the x-axis, while the y-axis shows the normalized (Z-score) gene expression. The dark phase of the 12h:12h light-dark cycle is represented in grey (dark phase begins at ZT12); **(D)** Heatmap showing the daily expression of all genes in the for-24h-nur-8h geneset, for nurses and foragers. Caste identity is indicated above the heatmap as a column annotation (red-foragers and blue-nurses). The for-24h-nur-8h geneset was clustered into four groups, and the cluster identity of each gene is indicated as row annotations (“cluster”). The majority of 8h-cycling genes in nurses, including the *Per* gene, belong to Cluster 1 and show a night-time peak in forager heads.

Having identified ultradian rhythms in gene expression, we asked if genes that oscillated in a circadian manner in forager brains, but not in nurses, were cycling in an ultradian manner in nurses. Indeed, we found that 325 (out of 2875) genes that cycled every 24h in foragers were not arrhythmic in nurses but differentially rhythmic genes (DRGs) that showed robust 8h (291 genes) or 12h (34 genes) rhythms (“for-24h-nur-8h” and “for-24h-nur-12h”, respectively; Fig. 5B). Remarkably, several components of the insect clock were among the 291 DRGs that cycled every 24h in foragers and every 8 hours in nurses: *Period* (*Per*)*, Shaggy* (*Sgg*; *Gsk3b* in mammals)*, Protein phosphatase 1b* (*Pp1b*), and *Protein phosphatase 1 at 13C* (*Pp1-13c* or *Pp1*) (Fig. 4, Table 1). This suggests that gene expression in nurse ant brains is, perhaps, not as arrhythmic as previously reported [28]. Instead, certain clock components in nurses seem to be cycling at a different harmonic compared to foragers, which could be partly facilitating the swift behavioral caste changes between foragers and nurses that have been observed in other studies [25, 48, 102]. As such, we continued our investigation into the genes that cycled every 24h in foragers and every 8h in nurses by asking if these DRGs play putative functional roles in regulating known clock-controlled processes as well as behavioral plasticity in ants.

### Chronobiological plasticity and behavioral division of labor in ants

In *Drosophila*, the circadian clock regulates daily rhythms in transcription via rhythmic binding of CLK and RNA Polymerase (Pol) II to the promoters of clock genes including *Per*, *Doubletime (Dbt; Ck1* in mammals) and *Shaggy (Sgg*, *Gsk3b* in mammals) [36, 103]. The kinase SGG regulates nuclear accumulation of the PER/TIM repressor complex [93, 104, 105], whereas DBT regulates its stability [106–108]. In addition to DBT, several other kinases (e.g., NEMO, CK2, and PKA) [106, 109–111] and a few phosphatases (e.g., PP1 and PP2a) [112, 113] have been identified as regulators of PER and PER/TIM stability in *Drosophila*. In our study, the daily changes in the expression of *Sgg*, *Dbt*, *Nemo*, *Pp1b* and *Pp1* mirrored the differentially rhythmic expression patterns of *Per* in the two ant castes (Fig. 5C, Table 1). Even though the 8h rhythms of *Dbt* (p=0.11) and *Nemo* (p=0.11) in nurse brains were not statistically significant, their expression patterns showed a strong phase coherence with *Per* (Additional File 5). Taken together, these findings further suggest that oscillations of key clock components at the third harmonic of the circadian rhythm in nurse brains might underlie a differentially regulated, yet functional, TTFL in this caste. Having core clock components that simply cycle at a different harmonic, versus not showing any rhythmicity at all, could indeed explain the ability of “around-the-clock” nurses to rapidly develop forager-like circadian activity, in behavior and gene expression, when their social context changes [25, 48, 102].

In the fruit fly *Drosophila,* the expression of *Per* and several other clock and clock-controlled genes peak during the night-time [103]. Similar to *Drosophila,* we observed a night-time peak in *Per* expression for *C. floridanus* foragers, which is also consistent with previous findings in fire ants and honeybees [14, 102]. Furthermore, hierarchical clustering of the DRGs that cycled every 24h in foragers and every 8h in nurses revealed that most of these DRGs clustered with *Per* (i.e., largely in-phase with the expression pattern of *Per* in foragers and nurses) (Fig. 5D, Additional File 7, sheet 1). Therefore, we hypothesized that the DRG-cluster in nurses that oscillated every 8h with a phase similar to *Period* would be enriched for some of the same biological processes performed by 24h cycling genes in foragers discussed above. Indeed, we found that the *Per*-like DRG-cluster was significantly enriched in functional annotations that we also identified in the night-peaking circadian gene cluster of foragers; the GO terms: transcriptional regulation (DNA-templated), transcriptional regulation by RNA Pol II, protein phosphorylation and GPCR signal transduction, (Additional File 7, sheet 2).

Moreover, the *Per*-like DRG cluster contained the muscarinic acetylcholine receptor gene *mAchR* and the insect dopamine/ecdysteroid receptor *DopEcR*, which have both been found to be clock-controlled in *Drosophila* [55, 114, 115]. The *mAchR* gene has a putative role in olfactory and mechanosensory signal transduction [116, 117]. Therefore, its differential clock-controlled regulation in foragers and nurses could be contributing to caste-specific behavioral phenotypes (Fig. 4). The same could be true for *DopEcR*, which modulates insect behavior by responding to dopamine, ecdysone and 20-hydroxyedysone [118–121]. In fact, dopamine is a known regulator of foraging activity in ants (reviewed in [122, 123]) and dopamine signaling has been found to be important in entraining the insect circadian clock as well as mediating clock-controlled behavioral phenotypes such as locomotion [124–126]. Moreover, studies in mammals suggest that certain dopaminergic oscillators are highly tunable and capable of generating ultradian rhythms in locomotor activity, independent of the circadian clock [127]. As such, our finding that forager and nurse ants respectively exhibit circadian and ultradian oscillations in the expression of genes that affect behavioral outputs, suggests that mechanistic links between chronobiological and behavioral plasticity in ants exist (Fig. 4).

It is not clear if the 8h rhythms in ant brain gene expression are endogenously produced or socially regulated, and what the functional aspects of such rhythms are, if any. However, the social insect literature does point to one likely role for the ability of nurses to track 8h periods: brood translocation. Workers of the carpenter ant species *Camponotus mus* have been observed to show daily rhythms in brood translocation behavior to move their brood between different temperature conditions. The measured time between the two daily brood translocations was exactly 8 hours [11, 128, 129]. This suggests that the 24h rhythm in thermal preference in *C. mus* nurses could be coupled with an 8h oscillator that drives the observed daily timing of temperature-dependent brood translocation. Brood translocation is important for larval development, and hence, has implications for colony fitness [12]. As such, 8h rhythms in behavioral outputs could have important adaptive functions. To begin to understand the potential roles for ultradian rhythms in the functioning of ant colonies, behavioral and molecular studies aimed at linking 8h transcriptional rhythms and brood translocation could provide a good first step.

### Plasticity in circadian entrainment and behavioral output pathways

The blue-light sensitive gene *cryptochrome*, which is essential for flies to entrain to light-dark cycles, is absent in both mammals and Hymenoptera [14, 130, 131]. As such, previous studies have suggested that the circadian clock in ants and honeybees likely resembles that of mammals, at least more so than the *Drosophila* clock does [14, 131]. However, not much is known about the molecular pathways that underlie circadian entrainment in ants. To assess the possibility of a mammalian-like entrainment pathway in ants, we queried the *C. floridanus* genome for orthologs of mammalian genes known to be involved in the circadian entrainment pathway (KEGG pathway: hsa04713). We found that *C. floridanus* possess one-to-one orthologs for most of the components in a mammalian-like entrainment pathway (Additional File 8), including the melatonin pathway that underlies dusk/dawn entrainment [78, 80]. Melatonin titers in the heads of adult honeybees show daily rhythms with crepuscular peaks at dusk and dawn [132]. As such, Hymenopterans might indeed have a melatonin-based entrainment pathway. Caste-specific differences in melatonin levels were also found, with lower titers in the heads of young nurse bees compared to older foraging individuals [132]. Our behavioral experiments indicated that nocturnal *C. floridanus* foragers are likely dusk-entrained. This suggests that a mammalian-like melatonin pathway might be involved in the dusk-entrainment of ant clocks as well. Indeed, the *C. floridanus* genome contains orthologs of both mammalian melatonin receptors – *trapped in endoderm 1* (*Tre1; MT1* in mammals) and *moody* (*MT2* in mammals) (Table 1, Fig. 4). While the gene expression of *moody* did not oscillate in either caste, *Tre1* expression showed circadian oscillation in foragers with a peak around dusk. In mammals, the activation of melatonin receptors at dusk or dawn triggers resetting of the clock through a signaling cascade that activates the kinase PKC [78]. Even though neither of the melatonin receptors were cycling in nurse brains, we found that *Pkc* oscillated every 8 hours while it does so every 24 hours in foragers (Table 1). Therefore, our data suggests that 24h rhythms in foragers could rely on a dusk entrainment pathway that likely involves melatonin-PKC signaling, while the 8h oscillatory rhythms in the nurse transcriptome could potentially be the result of an alternate, yet functional, *Pkc* activation pathway. Under the dark nest conditions in which nurses reside, this alternate 8h-oscillatory *Pkc* activation could be the main pathway to reset the clock, while this pathway gets overridden in foragers that experience light-dark cues and, thus, produce melatonin (Fig. 4).

Entrainment to external stimuli is central to the adaptive timing of rhythmic clock-controlled outputs such as extranidal foraging visits. While foragers receive both light and social cues, nurses primarily rely on social stimuli, which are likely to be different from those that foragers receive. As argued with regards to the melatonin-PKC signaling pathway, these entrainment cue differences might explain, at least partially, the differences in the clock-controlled output in the two ant castes. The differences that we found in clock-controlled gene expression could be giving rise to the observed presence of robust circadian activity in foragers and the absence of such rhythms in nurses. One such gene could be the *Foraging* (*For*) gene, which is known to regulate extranidal foraging activity of insects including ants [133–135]. Therefore, we hypothesized that the expression of *For* in forager ants would show rhythmic oscillations that mimic the daily foraging activity of the colony. Visual inspection of *For* expression patterns indicated a trend in rhythmicity that resembled the expression of *Period* in both foragers and nurses (Additional File 5). However, our eJTK analyses did not find any significance for these supposed gene expression trends in forager brains (tau = 24h, p = 0.1). As such, our hypothesis was not confirmed, which could be due to our experimental and analytical limitations. Alternatively, *For* might simply not be clock-controlled and rhythmic locomotory activity might be regulated by genes such as *Slowpoke* (*Slo*), a cycling potassium channel, which functions as a central regulator of rhythmic locomotion activity in flies [136, 137]. Indeed, *C. floridanus* contains a homolog of *Slo*, which appeared to be cycling every 24h in foragers (Fig. 4, Additional File 3, sheet 1). However, we did not find significant gene expression oscillations in nurses, though an 8h rhythmicity trend appeared to be there (tau = 8h, p = 0.17) (Additional File 5).

In addition to genes involved in locomotion and foraging, clock-controlled genes coding for neuropeptides and their receptors could be involved in regulating differentially rhythmic activity patterns in foragers and nurses. In flies, rhythmic activity patterns in total darkness have been related to the signaling pathway mediated by the neuropeptide Pigment Dispersing Factor (PDF) [36, 138–141]. PDF binds to the PDF receptor (PDFR) and triggers a signal transduction that increases cAMP levels and activates the protein kinase PKA [111]. A deficiency in PKA resulted in loss of fly locomotory rhythms even when *Per* oscillation was intact [142]. Moreover, PDF plays a central role in circadian timekeeping by mediating light input to the circadian clock neurons in the brain, coordinating pacemaker interactions among neurons, regulating the amplitude, period, and phase of circadian oscillations, and mediating output from the clock to other parts in the central brain [143–151]. Neurons that express PDF are present in the *C. floridanus* brain as well and could be mediating time-of-day information to brain regions involved in activity rhythms [68, 152–154]. In line with this, we found robust circadian rhythms in *Pdf, Pdfr* and *Pka* gene expression in the brains of *C. floridanus* foragers (Fig. 4, Table 1). However, nurse ants, which generally reside in dark nest chambers and demonstrate a lack of circadian rhythms in locomotion, did not exhibit circadian nor ultradian rhythmicity in *Pdf*, *Pdfr* and *Pka* expression (Fig. 4, Table 1). The absence of circadian locomotory rhythms in nurse ants could, thus, also be the result of a non-oscillatory PDF signaling pathway.

### Clock-control of differentially expressed genes

Past research has identified several genes and pathways that could be underlying behavioral division of labor [75, 155–159]. However, the extent of clock control over these key elements has not been explored yet. As such, we identified genes that were differentially expressed between the two ant castes throughout the day and determined if these differentially expressed genes (DEGs) showed any circadian or ultradian oscillations. Of the 10,038 expressed genes in the brains of *C. floridanus*, only 81 were significantly differentially expressed between the two behavioral groups throughout the day (fold change ≥ 2, q-value < 0.05; Additional File 9, sheet 1). Of these DEGs, 34 were significantly higher expressed in forager brains, and the remaining 47 were higher expressed in nurses (Fig. 6; Additional File 9, sheet 1). The 34 genes that were higher expressed in foragers comprised of several genes with unidentified functions and did not contain any significantly enriched GO terms. In contrast, the 47 genes that were higher expressed in nurses contained five maltase and five alpha-amylase genes which resulted in a significant enrichment for the GO terms carbohydrate metabolic process and catalytic activity (Additional File 9, sheet 2). This suggests that nurses might be metabolically more active than foragers, which is in line with previous findings from another ant species [75].

**Figure 6.**
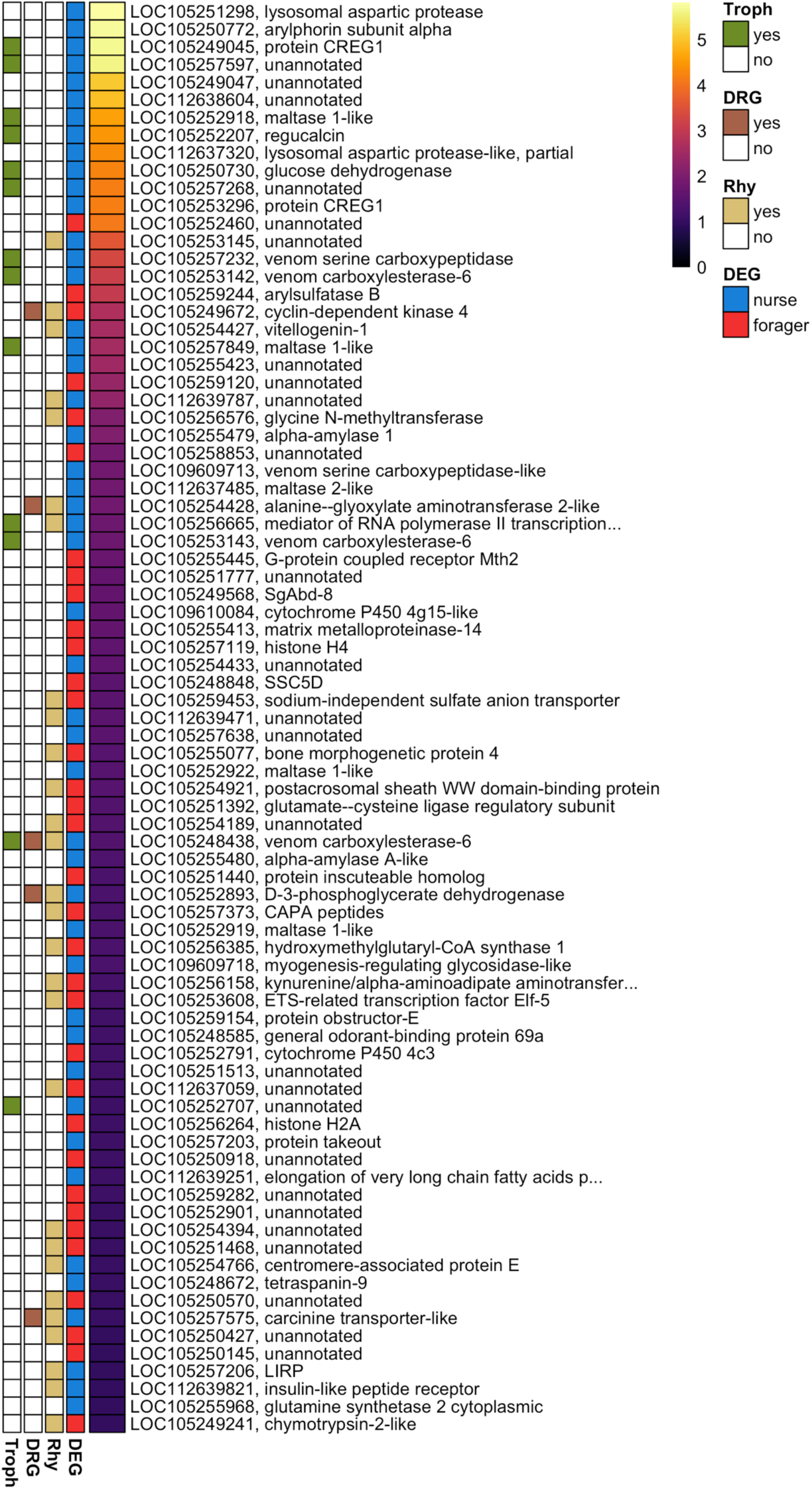
Differentially expressed genes between forager and nurse ant brains. Heatmap showing absolute (abs) log2-Fold-Change (log_2_FC) values for all 81 DEGs (q < 0.05 and abs(log_2_FC) ≥ 1), ordered from highest to lowest fold-change. The DEG column indicates if the gene is significantly higher expressed in foragers (red) or nurses (blue). For each DEG, the *C. floridanus* gene numbers and their blast annotations are provided. Genes with no blast annotation or annotated as uncharacterized protein are indicated as “unannotated”. The Rhy (rhythmic) column indicates genes that are significantly rhythmic in at least one of the ant castes. The DRG column indicates genes that are significantly rhythmic in both castes but oscillating at different periodicities. Genes that code for proteins previously found in the trophallactic fluid of *C. floridanus* are indicated in the Troph column.

Looking for oscillating genes among the DEGs that we identified in *C. floridanus*, we found that more than one-third (i.e., 28 of the 81 DEGs) were expressed rhythmically in either forager or nurse brains (Fig. 6). Of these clock-controlled DEGs, five genes oscillated at different periodicities in the two ant castes, providing further support for potential links between chronobiological and behavioral plasticity in *C. floridanus*. One of these differentially rhythmic genes, *Cyclin-dependent kinase 4* (*Cdk4*), was higher expressed and cycled every 12h in forager brains, while it cycled with an overall lower expression in nurse brains every 24h (Fig. 6, Additional File 9, sheet 1). Conserved in flies and mammals, *Cdk4* encodes a protein that regulates the JAK-STAT and TORC1 signaling pathway, and as such, is important for innate immune response and insulin signaling in insects. The Insulin/IGF-1 signaling (IIS) pathway is a known modulator of the circadian clock in insects [160], and a key pathway involved in longevity, fertility and behavioral division of labor in ants [161–163]. As such, the finding that *Cdk4* is both expressed at different levels throughout the day and cycling differently in foragers and nurses could be indicating a direct link between clock-controlled gene expression and division of labor (Fig. 4).

The other four differentially rhythmic DEGs, *Alanine—glyoxylate aminotransferase 2-like* (*Agxt2*), *D-3-phosphoglycerate dehydrogenase* (*Phgdh*), *Carcinine transporter-like* (*CarT*) and *Venom carboxylesterase-6*, showed a higher overall expression in nurse brains where they cycled every 24h, while foragers exhibited an 8h oscillation in expression (Fig. 6, Additional File 9, sheet 1). The gene *Agxt2* regulates nitric oxide (NO) signaling [164], which in *Drosophila* has been shown to mediate neuro-glial interactions that shape circadian locomotory rhythms [165, 166]. Additionally, there is a growing body of literature that suggests a key role of glia in maintaining circadian rhythms in activity and rest [166]. As such, glia could also be playing a role in regulating plasticity of locomotory rhythms in ants. However, *Agxt2* does not appear to be rhythmically expressed in *Drosophila* while we found it to be differentially rhythmic across behavioral ant castes. As such, its role in ants might be different (Fig. 4).

Like vertebrates, insects have functional N-methyl-D-aspartate (NMDA) glutamate receptors. Such receptors are thought to play a role in synaptic plasticity, memory, and neuronal development in vertebrates and mediate juvenile hormone (JH) biosynthesis in insects [167]. The gene *Phgdh* catalyzes the first step in L-serine synthesis [168, 169], thus, regulating the availability of L-serine, which controls NMDA receptor function [168, 170–172]. Therefore, *Phgdh* could be indirectly affecting JH levels in insects such as ants and result in different behaviors across the forager and nurse castes that appear to express this gene both in different quantities and at different oscillation rates (Fig. 4). This could be in conjunction with the differentially oscillating DEG *venomcarboxylesterase-6* (Fig. 6, Additional File 9, sheet 1), which modulates JH levels through its function as a JH esterase [73, 173] (more about this gene in the section below). Additionally, glutamate and its receptors have been found to regulate task specialization associated with division of labor in social insects [174–176], and expression of social traits in vertebrates including humans [177, 178]. Therefore, significant differences in *Phgdh*-mediated glutamate signaling in nurse and forager brains might be contributing to the caste-associated differences in social behavior (Fig. 4). Alternatively, *Phgdh* could be indirectly involved in photo-induced locomotor rhythms since glutamatergic neurotransmission is said to be important for circadian photoentrainment in both vertebrates and invertebrates [179, 180].

The differentially rhythmic gene *CarT* (Fig. 6, Additional File 9, sheet 1) also has an indirect role in phototransduction, but via histamine recycling [181]. The biogenic amine histamine is synthesized in photoreceptors and released from the compound eyes [182, 183] to regulate sleep via the (visual) photic and the (non-visual) motion detection pathways [184–187]. Eventually, the epithelial glia cells communicate with photoreceptor cells to drive conversion of histamine to carcinine [181]. CarT transports this carcinine back into photoreceptor neurons, which is an essential step in the histamine-carcinine cycle [181]. As such, overall different expression levels and oscillation patterns in *CarT* between nurses and foragers, possibly indirectly driven by different external photic and motion detection cues that induce histamine production, could be involved in regulating the different activity patterns of the two castes (Fig. 4). This differentially rhythmic DEG, as well as the other three we discussed above, can be indirectly tied to clock entrainment and clock-controlled locomotory output, which highlights the potential link between differential gene expression, circadian plasticity and behavioral plasticity in ants (Fig. 4). However, functional testing will be required to confirm and understand their exact roles.

Even though not much is known about the role of circadian clocks in regulating behavioral plasticity in ants, studies have looked into the molecular basis of behavioral division of labor and, in doing so, have identified several genes that seem to be central regulators of behavioral plasticity in social insects [188]. Caste-specific differences in larval storage proteins, especially Vitellogenin (Vg) and Arylphorin subunit alpha, and JH have been consistently found across social insects. In bees, for example, high *Vg* levels and low JH titers correlate with nurse-like behaviors [189], whereas downregulation of *Vg* results in increased JH titers and a behavioral state characteristic of forager bees [190]. In line with this, nurses of the fire ant *Solenopsis invicta* show significantly higher *Arylphorin subunit alpha* expression as compared to the foragers [191]. Consistent with these previous findings, we found *C. floridanus* nurse brains to have significantly higher *Arylphorin-subunit-alpha* (50-fold) and *Vg* (6-fold) expression as compared to foragers (Fig. 6, Additional File 9, sheet 1). Additionally, our data showed that *Vg* expression is significantly oscillating every 24h in nurse brains. Although not significant, *Arylphorin subunit alpha* also showed a *Vg*-like oscillatory expression in nurse brains (tau = 24h, p = 0.09) (Additional File 5). However, forager brains showed no such rhythms in *Vg* or *Arylphorin subunit alpha* expression. As such, our study provides further support for a role of *Vg* and *Arylphorin subunit alpha* in behavioral division of labor and highlights a putative clock-control of these genes in nurse brains (Fig. 4). Nevertheless, the potential functional role of a rhythmic *Vg* expression in ant physiology or behavior remains to be explored.

### Social regulation of differentially expressed genes

In addition to identifying several putatively clock-controlled DEGs, we found evidence for trophallactic fluid having a potential role in regulating division of labor in *C. floridanus.* Ants use trophallactic fluid as a way to exchange food and social cues. Such inter-individual interactions are usually more frequent within a behavioral caste [192]. This makes it likely that the trophallactic fluid contents differ between forgers and nurses, which might help maintain behavioral and physiological states associated with the specific castes. Nevertheless, a role for trophallactic fluid in regulating caste-specific gene expression has not been investigated yet. We do not currently know how and if the trophallactic fluid of forager and nurse ants differs. However, the trophallactic fluid of *C. floridanus* has been characterized by pooling fluid from both foragers and nurses [73]. We compared our list of DEGs between foragers and nurses with the previously reported proteins found in the trophallactic fluid of *C. floridanus* to investigate if trophallactic fluid could be a potential social regulator of caste-associated behavior. We found that more than a quarter (13 out of 47) of all genes that were significantly higher expressed in nurses compared to foragers encoded such orally transferred proteins (Fig. 6). Among these thirteen genes, only two showed significant daily oscillations in gene expression; the previously mentioned *venom-carboxylesterase-6* and a *mediator of RNA polymerase II transcription subunit alpha 12*.

Previous work showed that the trophallactic fluid of *C. floridanus* contained venom-carboxylesterase-6 JH esterases (JHEs). JHEs are enzymes that degrade JH in insect hemolymph, thus, regulating JH titers and its associated behaviors in ants [73, 173]. For instance, increasing JH levels in leafcutter ants results in increased phototaxis and extranidal activity [193]. The role of JH in regulating extranidal foraging has been demonstrated in honeybees as well [194, 195]. Moreover, experimentally introducing inhibitors of JHE, or JH itself, in the trophallactic network of *C. floridanus* workers increased larval growth and rate of pupation [69, 73]. In addition to affecting larval development, JHE levels were also shown to be affected by social context as the amount of venom-carboxylesterase-6 was significantly reduced in the trophallactic fluid of ants upon social isolation [73]. We found that all three copies of the putative JHE *venom-carboxylesterase-6* in the *C. floridanus* genome were significantly higher expressed in nurse brains, as compared to foragers (Fig. 6). This is in line with the expectation that nurses would have lower JH levels. The significantly higher expression levels of *venom-carboxylesterase-6* in nurses likely have a suppressing effect on JH-mediated foraging through the degradation of JH [194]. Additionally, as we mentioned above, the expression of *venom-carboxylesterase-6* showed differential rhythmicity between the two ant castes, oscillating every 24h in nurses and every 8h in foragers. While this was only the case for one of the three esterases, this indicates that there could be potential daily fluctuations of JH titers in in the nurse caste. The peak of the oscillating *venom-carboxylesterase-6* gene in nurse brains was around ZT12-14, which corresponds to the peak time of colony foraging that we found in *C. floridanus* (Fig. 1, Additional File 4 and 6). The *venom-carboxylesterase-6* mediated dip in JH levels could, thus, be contributing to a lower propensity of nurses to engage in extranidal tasks during peak colony foraging hours. In line with this reasoning, we found that the lowest dip in forager *venom-carboxylesterase-6* expression, and likely corresponding increased levels of JH, occur at ZT12, the onset of peak foraging activity (Fig. 1, Additional File 4 and 6). As such, the significantly different levels of JHEs, and therefore JH, in the trophallactic fluid of foragers and nurses, and the clock-controlled fluctuations within those expression levels, could be contributing to the regulation of differential locomotory activity in these castes (Fig. 4).

## Conclusion

The study presented here is providing a first look at the clock-controlled pathways in ants that could underlie caste-associated behavioral plasticity and sheds new light on the links between molecular timekeeping and behavioral division of labor in social insects. Understanding how an ant’s biological clock can predictably interact with its environment to produce distinct, yet stable, caste-associated chronotypes, lays the foundation for further molecular investigations into the role of biological clocks in regulating polyphenism in ant societies.

To produce high-interval time course data that reflects the transcriptional differences between forager and nurse ants throughout a 24h day, we used a behavioral setup that allowed us to reliably sample and obtain daily behavioral and brain gene expression patterns that appeared to be a good representation of these behavioral castes. In fact, the behavioral activity data that we were able to collect as part of these endeavors had high enough resolution to even identify how the colony is able to quickly get back on track with regards to food collection efforts after a disturbance. More importantly, we found a reduced circadian time keeping in nurses as compared to foragers. This was evidenced by the vastly different number of genes that oscillated every 24h in each ant caste, and the temporal segregation of clock-controlled processes, which is detectable in both castes but to a lesser extent in nurses. Our findings are, therefore, in line with the results of a previous study done in honeybees, which indicates that a difference in circadian gene repertoire between foragers and nurses could be a more general phenomenon within eusocial Hymenoptera, and likely contributes to the caste-specific differences observed in behavioral activity rhythms.

Moreover, many genes that showed a circadian expression in forager brains were expressed in an ultradian manner in nurses, instead of being entirely arrhythmic. Among the differentially rhythmic genes were essential components of the core and auxiliary feedback loops that form the endogenous clock of insects, as well as genes involved in metabolism, cellular communication and protein modification (Fig. 4). The ability of core clock and clock-controlled genes to oscillate at different harmonics of the circadian rhythm, and to switch oscillations from one periodicity to the other due to age or colony demands, might explain why chronotypes associated with ant behavioral castes are stable in undisturbed conditions, yet highly plastic and responsive to changes in their social context. However, it remains to be seen if the caste-associated differential rhythmicity that we observed is a general phenomenon across ant and other eusocial societies, or a species-specific trait. In addition, the potential for an actual adaptive function for maintaining both circadian and ultradian rhythms in ant colonies will have to be further explored.

In addition to the indications that caste-specific behavioral phenotypes could be the result of genes that oscillate at different speeds, we also found evidence that different functions of the same genes or pathways might be employed under the different environmental contexts that these ants are in. As such, the behavior of nurse ants that remain in a dark nest could be regulated by different functions of *Rho* or activation pathways of *Pkc* than foragers who are detecting light at set times of days, due to exposure to distinct set of social cues and colony environment (Fig. 4). Our enrichment analyses showed that foragers and nurses could be expressing different genes with similar functions during the subjective night-time, while they use the same genes during the subjective daytime. Additionally, we found evidence for a role of trophallactic fluid in regulating differential gene expression between foragers and nurses. Several of the differentially expressed genes showed robust daily rhythms in either forager or nurse brains, including *Vg* and *venom-carboxylesterase-6* that are known regulators of JH titers in insects (Fig. 4). Given the central role of Vg and JH in regulating division of labor in social insects, we propose that a mechanistic link between plasticity of the circadian clock and division of labor likely exists. Overall, this study allowed us to identify *C. floridanus* genes potentially involved in social entrainment of the endogenous clock, clock-controlled plasticity in behavior, and social regulation of division of labor.

## Methods

### Camponotus floridanus collection and husbandry

We collected a queen-absent colony of *C. floridanus* containing more than a thousand workers and abundant brood (eggs, larvae and pupa) from the University of Central Florida Arboretum in late April of 2019. We housed the colony in a fluon coated (BioQuip) plastic box (dimensions 42 x 29 cm, Rubbermaid) with a layer of damp plaster (Plaster of Paris) covering the bottom. We provided 15% sugar solution and water *ad libitum* and fed crickets to the colony every 2-3 days. We also provided the colony with multiple light-impervious, humid test-tube chambers (50 mL, Fisher Scientific) which they readily moved their brood into and used as a nest. Until the start of the experiment, we kept the colony in this setup inside a climate-controlled incubator (I36VL, Percival) at 25°C, 75% relative humidity (rH), and a 12h:12h light-dark (LD) cycle.

### Experimental setup and timeline

To allow for visible behavioral division of labor between morphologically indistinguishable forager and nurse ant castes (see definitions below), we built a formicarium consisting of a nest box and a foraging arena (42 x 29 cm each, Rubbermaid). Both boxes had a layer of damp plaster covering the bottom. We carved multiple grooves into the plaster of the nest box to imitate nest chambers and kept the box covered at all times to ensure completely dark conditions. We placed the nest in a temperature-controlled darkroom at constant temperature and humidity (25°C, 70% rH). The foraging arena was placed inside a climate-controlled incubator (I36VL, Percival) under a 12h:12h LD cycle without twilight cues. Lights ramped from zero to >2000 lux within a minute when lights were turned on at Zeitgeber Time, ZT24 (or, ZT0, which indicates the same time of day) and turned off within the same short time at ZT12 (Additional File 10). We maintained constant temperature (25°C) and humidity (75% rH) inside the incubator to ensure that the LD cycle was the primary rhythmically occurring cue, i.e., Zeitgeber, for circadian entrainment (Additional File 10). Abiotic factors in the foraging arena and nest box were monitored using HOBO data loggers (model U12, Onset) that logged light levels, temperature and humidity at 30 second intervals (Additional File 10). Food was provided *ad libitum* on an elevated circular feeding stage in the foraging arena to distinguish active feeding bouts from general extranidal visits (Additional File 11A). Feeders were replenished, and fresh frozen crickets were provided, every day between ZT2 and ZT4, throughout the experiment. The nest box was connected to the foraging arena with a 1.5m long plastic tube (i.e., *Tunnel*, Additional File 11A), which allowed ants to visit to the foraging arena at any time of the day.

Once the formicarium was set up, we transferred the entire colony along with brood into the foraging arena. To incentivize the colony to move their brood into the dark nest box, we kept the foraging arena under constant light for three consecutive days. This also aided in the resetting of their biological clocks to allow for synchronized entrainment to the 12h:12h LD cycle. After 5 days of initial entrainment, we identified and marked foragers for three consecutive days (Day 6-8, Fig. 1, see below for details on mark-and-recapture). This was followed by another four days of entrainment (Pre-sampling entrainment, Day 9-12, Fig. 1) before we sampled nurse and forager ants at two-hour intervals, spanning an entire LD cycle on day 13 (see below for sampling details).

### Colony activity monitoring

The extranidal or outside nest activity of the colony (called *activity* from here on) was used as a proxy for detecting rhythmicity in colony behavior. Before sampling ants for RNASeq, we analyzed the activity data to (a) confirm colony entrainment to the LD cycle, (b) identify peak activity hours for forager identification and painting, and (c) confirm pre-sampling entrainment after foragers had been marked. We monitored colony activity during the entire experimental period by recording time-lapse videos of the foraging arena using a modified infra-red enabled camera (GoPro Hero 6) at 4K resolution, set to capture one frame every 30 seconds at a wide field of view. To facilitate night-time recording, we installed a low intensity near-infrared light (850 nm, CMVision YY-IR30) above the foraging arena. We quantified extranidal activity throughout the experiment by counting the number of ants in the foraging arena on the feeding stage (FS) and off the feeding stage (FA, i.e., foraging arena) at one-hour intervals. The activity data can be found in Additional File 12.

### Identification of Camponotus floridanus behavioral castes

To measure and compare their daily rhythms in gene expression, we set out to sample minor worker ants of the behaviorally distinct foraging and nursing castes. We defined foragers as individuals that perform outside-nest (extranidal) tasks, including foraging for food. To identify foragers, we used a mark and recapture strategy. For three consecutive nights (Day 6-8, Fig. 1), we collected ants from the foraging arena during peak hours of extranidal activity (ZT13 to ZT16) as well as during relative dawn (ZT23 to ZT24). We marked new captures with a dab of white paint (Testors Enamel Paint) on their abdomen. Recaptures were marked with a second dab of white paint on their thorax. After painting, the ants were released back into the foraging arena. Since peak foraging hours took place during the night-time, we installed a 660 nm red lightbulb (Byingo LED) in the darkroom and wore a red headlamp (Petzl Tikka) to provide us with enough visibility to perform the mark-recapture, while simultaneously minimally disturbing the ants. We identified and marked more than a hundred foragers at the end of the three-day forager identification phase (109 doubly marked, and 39 singly marked). Post forager identification, the whole colony was left undisturbed and allowed to recover from potential stress for four consecutive days of pre-sampling entrainment, prior to sampling ants for RNASeq.

We defined nurses as ants that predominantly stay inside the dark nest chambers (intranidal) and care for brood. As such, we identified nurses as unmarked individuals in the colony that were unlikely to have gone outside the nest and were in contact with the brood. To confirm that the bulk of brood care inside the nest was performed by unmarked ants, and not marked foragers, we performed qualitative intermittent behavioral observations for a total of 1-2 hours per day during the pre-sampling entrainment period that followed mark-recapture (Days 9-11, Fig. 1). We observed the nest chambers under the same red light (660 nm) that illuminated the darkroom. Monitoring behavior inside the nest confirmed that marked “foragers” were less likely to be in direct contact with the brood (i.e., walking on the brood pile or grooming brood) and were not seen to be involved in brood relocation within the nest chambers. As such, we identified nurses as “unmarked” individuals found in direct contact with the brood or involved in brood care including relocation.

### Ant sampling and brain dissections

After identifying foragers and nurses and 12 days of colony entrainment to the 12h:12h LD, we collected ants for RNASeq under the same light-dark regime. We sampled ants from the colony every 2 hours over a 24-hour period, starting two hours after lights were turned on (ZT2) (Additional File 11B). At each sampling time point, we collected three foragers and three nurses from the colony and transferred them into individually labelled cryotubes (USA Scientific) for immediate flash freezing in liquid nitrogen. The whole process, from collection to flash freezing, took less than 60 seconds per sampled ant. Since *C. floridanus* foraging activity is predominantly nocturnal, we sampled foragers from inside the dark nest box during the light phase, and from the foraging arena during the dark phase (Additional File 11B). Nurses were always collected from inside the nest box. For sampling under dark conditions, we used the same intensity red-light as described for the mark-recapture and behavioral observations described above. Using this sampling regime, we collected 72 ants, which were stored at −80°C until brain dissection.

To compare transcriptome-wide daily gene expression patterns in the brain tissue of foragers and nurses, we performed brain dissections of individual flash-frozen ants in ice-cold Hanks’ balanced salt solution (HBSS) buffer under a dissecting microscope. To further preserve RNA integrity and quality, we performed brain dissections as swiftly as possible: brain dissections of individual foragers took an average of 4.6 (±0.7) mins, whereas for a nurse it took 4.5 (±0.5) mins. Immediately after dissection, brains were flash frozen again in cryotubes (USA Scientific) kept on dry ice. For each behavioral caste, at each sampling time point, we pooled three individually dissected brain samples for RNA extraction and sequencing (Additional File 11C). The resulting 24 samples were again stored at −80°C until RNA extraction and library preparation. This sampling approach was designed to adhere to current recommendations for genome-wide time course studies using non-model systems [60, 196]. By pooling triplicates, we have accounted for intra-colony variation while still being able to choose a high sampling frequency (every 2h) and read depth per sample (≥ 20M per sample, see below) in order to maximize accurate detection of the majority of cycling transcripts in *C. floridanus* brains [197].

### RNA extraction, library preparation and RNASeq

To obtain time course transcriptomes for each of the behavioral castes, we extracted total RNA to prepare sequencing libraries for Illumina short-read sequencing. Two frozen steel ball bearings (5/32” type 2B, grade 300, Wheels Manufacturing) were added to each cryotube containing the pooled brain tissues to homogenize them using a 1600 MiniG tissue homogenizer (SPEX) at 1300 rpm for 30 sec while keeping the samples frozen. We isolated total RNA from the disrupted brain tissues with Trizol (Ambion) followed by a wash with chloroform (Sigma) and a purification step using RNeasy MinElute Cleanup columns and buffers (Qiagen) [198]. For each library preparation, we used 500 ng total RNA to extract mRNA with poly-A magnetic beads (NEB) and converted this mRNA to 280-300 bp cDNA fragments using the Ultra II Directional Kit (NEB). Unique sequencing adapters were added to each cDNA library for multiplexing (NEB). All twenty-four cDNA libraries were sequenced as 50 bp single-end reads using two lanes on an Illumina HiSeq1500 at the Laboratory for Functional Genome Analysis (Ludwig-Maximilians-Universitat Gene Center, Munich). Read data are available under BioProject PRJNA704762. After sequencing, we removed sequencing adapters and low-quality reads from our RNASeq data with BBDuk [199] as a plug-in in Geneious (parameters: right end-low quality trim, minimum 20; trim both ends - minimum length 25 bp) (Biomatters). Post-trimming, we retained an average of 22 million reads per sample, which is well beyond the minimal read depth sufficient to identify the majority of high amplitude circadian transcripts in insects (Li et al., 2014) [197]. Subsequently, we used HISAT2 [200] to map transcripts to the latest Cflo v7.5 genome [201], followed by normalizing each sample to Fragments Per Kilobase of transcript per Million (FPKM) with Cuffdiff [202].

### Data analyses

We confirmed daily rhythms in colony activity with the WaveletComp package [74]. Using wavelet analyses, we investigated the extranidal activity of foragers for the presence of circadian rhythms in colony behavior, the potential presence of ultradian rhythms, and to infer synchronicity between the number of ants actively feeding or present on the feeding stage (*FS*), and those present in the remainder of the foraging arena (*FA*).

We used the rhythmicity detection algorithm empirical JTK-Cycle (eJTK) [76, 77] to test for significant circadian and ultradian rhythms in gene expression in foragers and nurses using waveforms of period lengths (tau) equal to 24h, 12h and 8h. Only genes that had diel expression values ≥1 FPKM for at least half of all sampled timepoints were tested for rhythmicity. For a set period length, a gene was considered to be significantly rhythmic if it had a Gamma p-value < 0.05. To test if certain genes could be clustered together based on similar temporal peak activity, we used an agglomerative hierarchical clustering framework (method = complete linkage) using the ‘hclust’ function in the ‘stats’ package for R.

Time-course sampling of foragers and nurses enabled us to account for diel fluctuations in expression levels when identifying genes that were differentially expressed between the two ant groups throughout the day (i.e., DEGs). To determine differentially expressed genes, we used the linear modelling framework proposed in LimoRhyde [203], but without an interaction between treatment and time. A gene was considered differentially expressed if treatment was found to be a significant predictor (at 5% FDR) and the difference in mean diel expression between foragers and nurses was at least 2-fold (i.e., abs(log_2_-fold-change) ≥ 1). LimoRhyde is generally used to test if genes are differentially rhythmic in phase or amplitude, inferred from a significant interaction between treatment and time. However, we did not find significant differences in phase or amplitude for any of the circadian genes. Therefore, we indicated a gene as differentially rhythmic (i.e., DRGs) if it significantly cycled in both ant castes but with different period lengths.

To perform functional enrichment analyses of significant gene sets, we wrote a customized function that performs a hypergeometric test through the *dhyper* function in R. The code is available on GitHub (https://github.com/debekkerlab/Will_et_al_2020). The function takes the following inputs: (1) user-provided geneset to test enrichment on, (2) user-provided background geneset to test enrichment against, and (3) functional gene annotations (e.g., GO terms) to test enrichment for. Among other things, the function outputs a Benjamini Hochberg-corrected p-value for each annotation term to indicate if it is significantly enriched in the test geneset. We used all genes that were found to be “expressed” (≥ 1 FPKM expression for at least one sample) in the brains of foragers or nurses as the background geneset for functional enrichment tests. To analyze the functional enrichment of Gene Ontology (GO) predictions, we used the GO term annotations [71] for the most recent *C. floridanus* genome (v 7.5) [201]. We only tested terms annotated for at least 5 protein coding genes and significance was inferred at 5% FDR.

Homologs of known core-clock genes (*cgs*) and clock-modulator genes (*cmgs*) *in C. floridanus* were identified using previously published hidden-markov-models (HMMs) for well-characterized clock proteins of two model organisms: *Drosophila melanogaster* and *Mus musculus* [204]. We used *hmmersearch* to query these HMM profiles against the entire *C. floridanus* proteome (Cflo_v7.5) [201] with default parameters (HMMER v3.2.1 [205]). To identify orthologs shared between *C. floridanus* and flies, mammals or honey bees we used proteinortho5 [206].

All data wrangling, statistical tests and graphical visualizations were performed in RStudio [207] using the R programming language v3.5.1 [208]. Heatmaps were generated using the pheatmap [209] and viridis [210] packages. Upset diagrams were used to visualize intersecting gene sets using the UpsetR package [211].

## Declarations

### Ethics approval and consent to participate

Not applicable

### Consent for publication

Not applicable

### Availability of data and materials

Raw sequencing reads generated for this study have been deposited in NCBI under BioProject PRJNA704762. The datasets supporting the conclusions of this article are included within the article and its additional files. Data analysis and visualization for this study was done using code written in R, Python and Bash, and can be found through GitHub (https://github.com/debekkerlab/Das_et_al_2021). Additionally, an RSQLite database containing all processed data can be provided upon request.

## Funding

This work was supported by NSF Career Award 1941546 and start-up funds from the University of Central Florida made available to CdB.

### Author’s contributions

BD and CdB both conceived of the study, analyzed the data and have written the manuscript. All experiments were performed by BD.

## Acknowledgements

We thank the Laboratory for Functional Genome Analysis and Genomics Services Unit and Adnreas Brachmann at the Ludwig Maximilians Universität for sequencing support. We also thank Ian Will for his assistance in the processing of our raw sequencing data, Daniel A. Friedmann for sharing brain dissection videos, Thienthanh Trinh, Veronica Urgiles, Leo Ohyama and Jordan Dowell for extensive discussions about the study, and Alicia Formanack for providing feedback on early drafts of the manuscript.

## Additional Files

### Additional File 1

Ultradian rhythms in colony behavior. (**A)** The length of day (daylength in hours, top) and dusk (twilight in minutes, bottom) in Orlando, for every single day in 2019, is shown. Data was obtained from www.timeanddate.com. The vertical grey line indicates the time-of-year (May, 2019) during which the *C. floridanus* colony was collected and all experiments were performed. (**B)** Recreated activity profiles of feeding bouts (shown in red lines) using decomposed 24h, 12h, and both (12h+24h) waveforms plotted on top of the observed activity data (black lines). The vertical dashed lines indicate the time period during which the 12h rhythms were found to be significant. The x-axis shows the Cumulative ZT (in hours) since mark-and-recapture. The y-axis indicates colony feeding activity (FS) as the number of ants present on the feeding stage at a given time point. The 12h:12h light-dark cycles are indicated in white (lights on) and grey (lights off) at the top. (PNG, 620 KB)

### Additional File 2

General patterns of gene expression in forager and nurse ants. The excel file contains four worksheets. (**Sheet 1**) The excel worksheet contains list of genes that displayed “no expression” (FPKM = 0) and “low expression” (0 < FPKM < 1) in the brains of *C. floridanus* foragers and nurses. In addition to the gene symbols (column: gene_name), the blast annotation (column: blast_annotation) and expression data for foragers (column: X2F to X24F) and nurses (column: X2N to X24N) are also provided. (**Sheet 2**) Results of GO enrichment analyses for genes that show “no expression” and “low expression”. (**Sheet 3**) List of genes that are expressed only in forager brains or nurse brains. (**Sheet 4**) Results of GO enrichment analyses for genes expressed only in forager brains or nurse brains. (XLS, 1.7 MB)

### Additional File 3

Circadian gene expression in forager and nurse brains. The excel file contains five worksheets. (**Sheet 1**) eTJK output for all tested genes in forager brains, including their gene number and normalized expression levels for each time point, sorted based on significance. (**Sheet 2**) eTJK output for all tested genes in nurse brains, including their gene number and normalized expression levels for each time point, sorted based on significance. (**Sheet 3**) List of genes that show significant 24h rhythms in forager brains, their cluster identity (corresponding to Fig. 3B), and normalized gene expression for all forager samples. (**Sheet 4**) List of genes that show significant 24h rhythms in nurse brains, their cluster identity (corresponding to Fig. 3C), and normalized gene expression for all nurse samples. (**Sheet 5**) GO enrichment results for circadian genes in foragers (for-24h) and nurses (nur-24h) that peak during the day (day-peaking clusters) and night (night-peaking clusters). Also includes the enrichment results for day-peaking cluster of overlapping for-24h and nur-24h genes (for-24h-nur-24h). (XLS, 12.6 MB)

### Additional File 4

Core clock and clock-controlled genes in *C. floridanus*. The excel worksheet contains the list of fly- and mammalian-like core clock genes and clock-controlled genes identified in *C. floridanus*, along with the hmmersearch results. (XLS, 39 KB)

### Additional File 5

The figure shows the expression patterns of several genes with a rhythmic trend that are discussed in the text. The forager expression is shown in red and nurse expression in blue. For each gene, the periodicity of rhythmic expression tested in forager and nurse brains are shown along with p-values obtained from eJTK (in parenthesis). The y-axis shows gene expression (z-score) and the x-axis shows the Zeitgeber Time (in hours). Dark phase of the 12h:12h light-dark cycle is represented in grey (dark phase begins at ZT12). (PNG, 1.1 MB)

### Additional File 6

Ultradian gene expression in forager and nurse brains. The excel file contains four worksheets. (**Sheet 1**) Results of eTJK testing for significant 12h periodicity in forager brain gene expression. (**Sheet 2**) Results of eTJK testing for significant 12h periodicity in nurse brain gene expression. (**Sheet 3**) Results of eTJK testing for significant 8h periodicity in forager brain gene expression. (**Sheet 4**) Results of eTJK testing for significant 8h periodicity in nurse brain gene expression. (XLS, 14.1 MB)

### Additional File 7

Differentially rhythmic genes. The excel file contains two worksheets. (**Sheet 1**) List of genes that cycle every 24h in forager brains but every 8h in nurses (for-24h-nur-8h). In addition to gene symbol, gene annotations and normalized expression, the cluster identity of each gene upon hierarchical clustering is provided. (**Sheet 2**) GO enrichment results for “for-24h-nur-8h” genes belonging to cluster 1 that also contains the *Per* gene. (XLS, 251 KB)

### Additional File 8

Components of mammalian circadian entrainment pathway. The excel worksheet contains a list of mammalian genes that are involved in circadian entrainment pathway (KEGG pathway: hsa04713) and their orthologs in *C. floridanus* (if present). Additionally, the annotation, results from eJTK and normalized expression of *C. floridanus* orthologs is provided. (XLS, 48 KB)

### Additional File 9

Genes differently expressed between forager and nurse brains. The excel file contains three worksheets. (**Sheet 1**) LimoRhyde results for all genes tested for differential gene expression. (**Sheet 2**) GO enrichment results for genes significantly higher expressed in nurse brains as compared to forager brains. (XLS, 1.9 MB)

### Additional File 10

Abiotic conditions in the experimental foraging arena and the nest box. The figure shows the data for light intensity, temperature and humidity in the foraging arena and the nest box of the experimental setup, collected using HOBO data loggers. Data is shown for Day 12 and Day 13 of the experiment. (PNG, 600 KB)

### Additional File 11

Ant colony setup and experimental design. (**A**) The figure shows the ant colony setup used for the experiment. The scheme for sampling ants from the colony is shown in (**B**) and several of the key steps from sampling to RNASeq are shown in (**C**). (PNG, 777 KB)

### Additional File 12

Colony foraging and feeding activity data. The excel worksheet contains the number of ants observed on the feeding stage (FS) and involved in general foraging activity (FA). Total activity (Total) was defined as the sum of FS and FA. Experimental phases: Initial entrainment (Entrain-I), Mark-and-recapture (Painting), Pre-sampling entrainment (Entrain-II), Sampling, and Post-sampling entrainment (Entrain-III). (CSV, 45 KB)

